# Incorporating FDR into the assessment of nanopore sequencing for the reliable detection of DNA modifications in real applications

**DOI:** 10.1101/2024.11.19.624260

**Authors:** Yimeng Kong, Hao Chen, Edward A. Mead, Yanchun Zhang, Christian E. Loo, Yu Fan, Mi Ni, Emma L. Thorn, Laura Zuluaga, Ketan K. Badani, Fanny Elahi, John Crary, Xue-Song Zhang, Rahul M. Kohli, Gang Fang

## Abstract

While nanopore sequencing is increasingly used for mapping DNA modifications, it is important to recognize associated false-positive calls, as they can mislead biological interpretations. To assist biologists and methods developers, we describe a framework, modFDR, for rigorous evaluation that emphasizes the use of the false discovery rate with rationally designed negative controls capturing both general background and confounding modifications. Our critical assessment across multiple DNA modifications shows that while nanopore sequencing performs reliably for high-abundance modifications—including 5-methylcytosine (5mC) at CpG sites in mammalian cells and 5-hydroxymethylcytosine (5hmC) in mammalian brain cells—it produces a substantial fraction of false-positive detections for low-abundance modifications, such as 5mC at CpH sites, 5hmC, and N6-methyldeoxyadenine (6mA) in most mammalian cell types. Although newer models improve certain aspects, systematic false positives remain, and we further observe elevated false negatives for 5mCpG when benchmarked against orthogonal enzymatic methods. This study highlights the urgent need to incorporate modFDR into future methods development, evaluation, and biological studies, and advocates prioritizing nanopore sequencing for mapping abundant rather than rare modifications in biomedical applications.

## Introduction

The advent of nanopore sequencing technology has revolutionized the landscape of genomic research^1,2^. Not only does the approach generate high throughput long reads, but it also has a distinctive ability for the direct mapping of DNA modifications^3^. By circumventing the conventional requirements for chemical, antibody, or enzymatic treatments, nanopore sequencing enables the analysis of native DNA^3,4^. The capacity for resolving both primary nucleotide sequence and DNA modifications along individual long reads greatly empowers the study of human epigenomes in health and diseases, as well as a broad spectrum of species^5–8^.

DNA modifications, such as 5-methylcytosine (5mC) and its derivative 5-hydroxymethylcytosine (5hmC), play a crucial role in the regulation of genomic functions and are pivotal to understanding the complex mechanisms underlying various biological processes and diseases^9–11^. However, as the adoption of nanopore sequencing expands and the repertoire of computational tools for detecting DNA modifications grows, it has become increasingly important to critically assess the technology’s fidelity, particularly to identify spurious or problematic modification calls. Misinterpretation of such modification calls can mislead further biological studies and clinical applications^4,12–15^.

A notable example was the debate over DNA N6-methyladenine (6mA) in the mammalian genome. In brief, 6mA was thought to be exclusively present in the DNA of bacteria until several studies reported its detection in nematodes, insects, mice, and humans^16–19^ and proposed important functions of 6mA in human development and diseases^19^. However, subsequent studies then discovered multiple confounding factors and raised caution about the level of 6mA in the human genome^14,15,20,21^, leading to an active debate. The debate was well clarified when bacterial contamination was demonstrated to have contributed to the significantly overestimated 6mA abundance in the mammalian genome^4,12^. This debate and confusion reflected a critical caveat when researchers might use an arbitrary cut-off to make 6mA calls from the mammalian genome without a rigorous false positive analysis, trying to be “consistent” with liquid chromatography with tandem mass spectrometry (LC-MS/MS) estimates ^4^. In another example of confusion caused by different sequencing modalities, several studies using nanopore sequencing and/or whole genome bisulfite sequencing reported extensive 5mC in mitochondrial genomes (mtDNA) across multiple species^22,23^, while other studies suggested that mtDNA methylation levels had been overestimated due to confounding factors^24–27^. This debate was addressed when subsequent studies performed rigorous method evaluation^25^ and reported the 5mC level is extremely low in mammalian mtDNA, calling into question the previously described functional roles of 5mC in mtDNA metabolism.

These examples of false positive detections highlight the critical need for rigorous development and evaluation of modification mapping methods to prevent erroneous assumptions that could misguide future research. Motivated to address these challenges, we recently wrote a perspective detailing the pitfalls of detecting DNA and RNA modifications and strategies to navigate them ^4^. One of the most important pitfalls that requires attention is the risk of false positives when mapping DNA modifications of low abundance. A method may perform very well in detecting DNA modification of high abundance in a genome but can give mostly false positive calls when applied to a sample with low abundance of the same modification. Essentially, this is related to the important statistical concept of false discovery rate (FDR). Fundamentally, the reliability of a modification mapping technology not only depends on the intrinsic properties of the technology itself but also the abundance of the DNA modification of interest in a specific sample. However, existing methods for modification detection are usually trained and evaluated using datasets with predefined ratios of modified to non-modified bases (e.g., 1:1) that do not represent physiologically relevant levels of these modifications (e.g., 5hmC and 6mA are much less abundant than 5mC in most human cell types). This gap between method development and their ultimate applications underlines the critical need for FDR estimation to safeguard data interpretation.

In this study, we used nanopore sequencing, one of the widely used long-read technologies for mapping a variety of DNA (and RNA) modifications, to showcase the problem of false positives and necessities of FDR evaluation. We generated data using the latest R10.4.1 nanopore sequencing kits and critically assessed several versions of the official software for modification calling including a very recent release (as of June 15, 2026), which was reported to have high accuracy for detecting 5mC, 5hmC, and 6mA. Our assessment revealed contrasting performance of these models based on the abundance of the modifications. The technology exhibits high reliability for detecting high-abundance modifications, such as 5mC at CpG sites in mammalian genomes. However, it demonstrates a marked propensity for false positives when detecting low-abundance modifications. For instance, we observed this phenomenon with 5mC at non-CpG sites (namely 5mCpH) and 5hmC, which, although common in mammalian brain cells, are considerably less prevalent in other cell types^28–32^. This disparity in detection reliability is also evident with 6mA, which is abundant in bacteria and certain protozoan species, but occurs at much lower levels in higher eukaryotes^14,15,20,21^.

To mitigate the risk of false positive calls and provide guidance for broad users and tool developers, we present a framework, modFDR, that highlights the use of rationally designed negative controls (capturing both general background and confounding modifications) and FDR analysis to evaluate the reliability of detected modification events. This approach aims to help researchers navigate through the pitfalls of nanopore sequencing, particularly in the detection of low-abundance modifications (**Fig. 1a**).

**Fig. 1.**
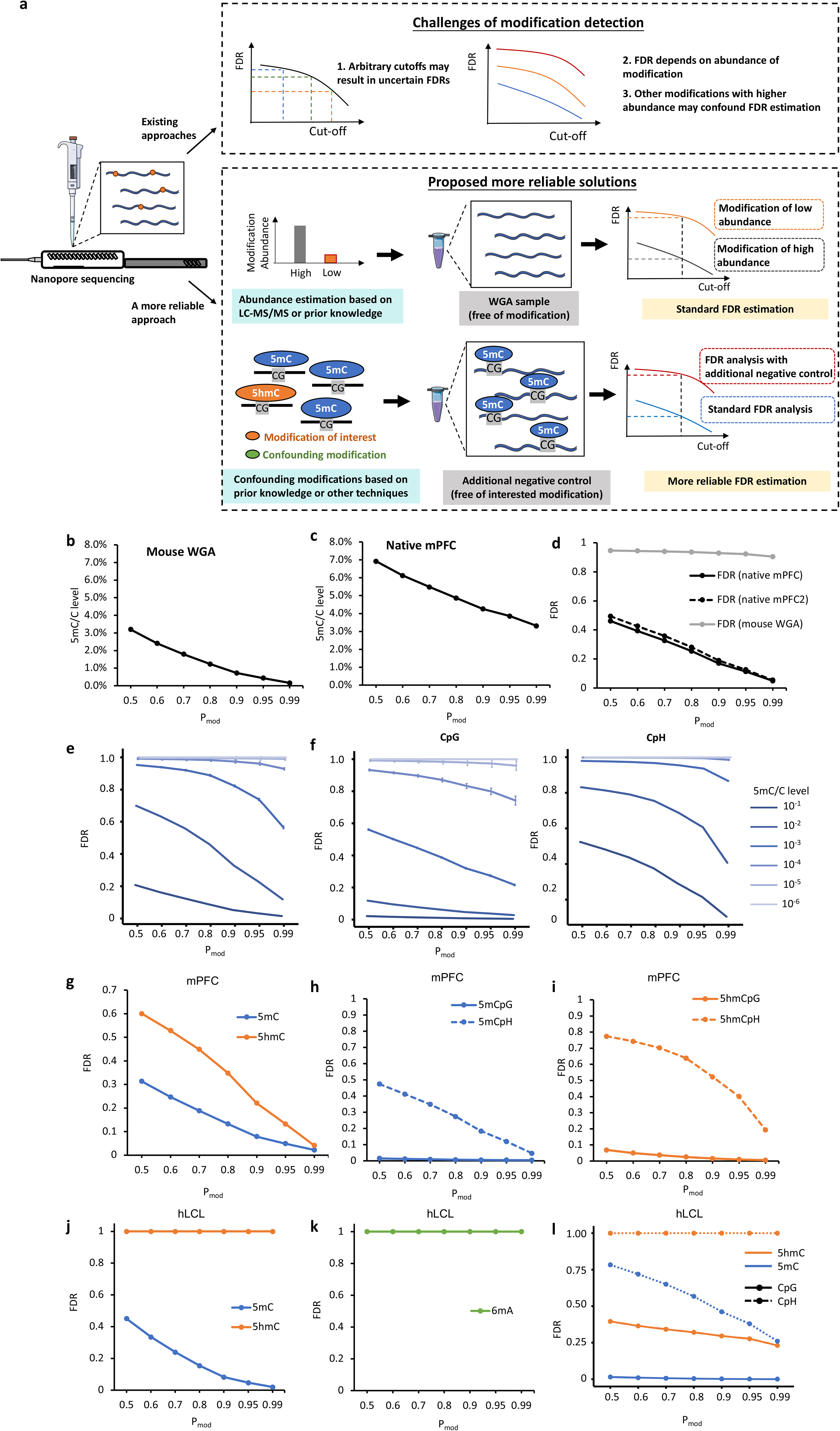
A framework for the rigorous evaluation of DNA modification detected by nanopore sequencing, and demonstration of false positives and the use of false discovery rate (FDR) to evaluate multiple forms of DNA modification called from mPFC and hLCL samples. a, The framework for evaluating the reliability of detected modifications using negative controls and FDRs. The use of rationally designed negative controls tailored based on prior knowledge including modification levels, genome contexts and coexistence of abundant confounding modifications. b, False positive 5mC sites among the total C (%) (*y-axis*) in mPFC gDNA subjected to whole genome amplification (WGA, free of modification) by applying different thresholds on modification probability (P_mod_) assigned by the official DORADO software (*x-axis*). Unmethylated C levels (%) for native mPFC and mouse WGA samples among all P_mod_ thresholds were shown in Supplementary Fig. 2. c, 5mC sites among the total C (%) (*y-axis*) identified on reads from native mPFC gDNA across different thresholds on P_mod_ (*x-axis*). d, FDR evaluation of read-level 5mC detection on mPFC sample (*y-axis*) across different thresholds on P_mod_ (*x-axis*). 2-round WGA on mPFC2 was compared the 1-round WGA on mPFC2 to estimate the FDR of the WGA sample (Methods). e, FDR evaluation for a list of samples that have 5mC/C levels with different orders of magnitude, which were simulated by randomly mixing native reads with WGA reads to mimic a number of 5mC/C levels (Methods). Three independent simulations were performed for each condition and error bars are standard deviation. f, FDR evaluation for 5mC in CpG sites (left) and CpH sites (right) in the simulated samples with varying 5mC/C levels across different magnitude orders (e). Three independent simulations were performed for each condition and error bars are standard deviation. g, FDR evaluation of 5mC and 5hmC calls made from the native mPFC sample using DORADO software (v0.5.3) with the 5mC_5hmC model (v4.3.0). h, FDR evaluation of 5mC made at CpG and CpH sites in the native mPFC sample using DORADO software (v0.5.3) with the 5mC_5hmC model (v4.3.0). i, FDR evaluation of 5hmC made at CpG and CpH sites in the native mPFC sample using DORADO software (v0.5.3) with the 5mC_5hmC model (v4.3.0). j, FDR evaluation for read-level 5mC and 5hmC calls from hLCL across different thresholds on P_mod_ (*x-axis*). k, FDR evaluation for read-level 6mA calls from hLCL. l, FDR evaluation of 5mC and 5hmC calls at CpG and CpH sites from hLCL.

## Results

### False positive calls can dominate detection when the DNA modification of interest has low abundance

To illustrate false positive calls, we first performed a whole genome amplification (WGA) of a mouse genomic DNA (gDNA) sample. The amplified DNA from the WGA process is essentially modification-free^4,33^ and exhibited fairly even amplification across the genome, serving as a negative control (**Supplementary Fig. 1**). The WGA sample was sequenced with the latest R10.4.1 kit and flow cell (**Supplementary Table 1**). Because many studies focused on 5mC methylation profiles in mammals, our study started from the read-level 5mC modification calling using the latest (as of June 15, 2026) two-state model (dna_r10.4.1_e8.2_400bps_sup@v4.2.0, hereafter v4.2.0) to differentiating 5mCs from Cs by official software DORADO (**Methods, Supplementary Table 2**). Although no or very low-level 5mC sites were expected from the WGA sample, 0.16%-3.19% of unmethylated C sites were called as 5mC from nanopore reads, representing false positives (**Fig. 1b; Supplementary Fig. 2**). The 3.19%-0.16% of false positive calls correspond to different thresholds on the read-level classification probability (P_mod_) in the DORADO output, ranging from low-confident (0.5) to high-confident (0.99) (**Fig. 1b; Supplementary Fig. 2; Methods**). As expected, the percentage of false positive 5mC sites decreased as the threshold increased. While 3.19% false positive 5mC sites were detected with a P_mod_ threshold of 0.5, ∼0.16% false positive 5mC sites were detected with a P_mod_ threshold of 0.99 (**Fig. 1b**). Consistent results were observed across various WGA preparation methods (**Methods**), validating that the WGA samples are reliable negative controls (**Supplementary Fig. 3**). This analysis demonstrated that even with a high confident threshold, the nanopore read-level 5mC calling by the two-state model can still make millions of false positive calls across a mammalian genome.

Next, we performed the same analysis on the native gDNA samples extracted from a mouse prefrontal cortex (mPFC). 6.92%-3.31% of unmethylated C sites were called as 5mCs across the same ranges (0.5-0.99 of P_mod_ thresholds) (**Fig. 1c; Supplementary Figs. 2 & 3**). While the percentage of unmethylated C sites changed dramatically with increased (more strict) P_mod_ threshold, the percentage of 5mC sites stays fairly stable (**Supplementary Fig. 2**), which is largely consistent with previously reported levels of 5mC/C (4-5% as estimated by LC-MS/MS) in mPFC^34^.

### The use of FDR to safeguard DNA modification detection across various abundance levels

Our above analysis demonstrates that the reliability of modification calls using the same calling method (DORADO in this case) depends on the abundance of a modification of interest in a specific sample. While a seemingly small proportion (0.16%) of false positive calls may seem negligible in the mPFC sample (with abundant 5mC), it could significantly mislead any biological interpretations in samples with the low 5mC/C levels, such as the WGA sample. This statistical concept is well established and is nicely captured by the measure FDR, defined as the ratio of N_fp_/N_p_, where N_p_ is the total number of all positive modification calls detected from the native sample (true positive N_tp_ + false positive calls N_fp_), and N_fp_ can be inferred from the negative WGA sample (free of modification) (**Supplementary Fig. 4**).

FDR decreases from 0.46 to 0.05 for the native mPFC gDNA data, as the P_mod_ threshold increased from 0.5 to 0.99, consistently with the FDR of a biological replicate (mPFC2) (**Fig. 1d**). To apply FDR analysis to the WGA dataset, we generated 2-round WGA dataset on mPFC2 to estimate N_fp_, and compared it with the 1-round WGA dataset of mPFC2 across different P_mod_ thresholds. Notably, the FDR remained >0.9 for the mouse WGA data across P_mod_ thresholds (**Fig. 1d**). Taking the P_mod_ threshold of 0.99 for example, 90.46% of 5mC events called from the mouse WGA data are false positives, while only 4.9% of the 5mC sites called from the native mPFC data are false positives.

The mouse WGA and native mPFC data described above represent two cases with either very low or high abundant 5mC. Although 5mC is generally abundant in mammalian genomes, much lower 5mC levels have been observed in other eukaryotic species such as yeast^35^ and Drosophila^35^, as well as in mammalian mtDNA^24–27^.

To further illustrate the use of FDR across various abundance levels of 5mC (representing its wide range across different eukaryotic and prokaryotic genomes), we simulated a series of 5mC/C levels across different orders of magnitude. Specifically, we randomly mixed nanopore reads from the native mPFC and mouse WGA samples at different proportions creating 5mC/C levels from 10^-1^ to 10^-6^. As shown in **Fig. 1e**, FDR increases as 5mC/C level decreases: for 5mC/C levels below 10^-3^, FDR stays close to 1, meaning the vast majority of the 5mC calls are false positives within a gDNA sample with rare 5mCs. This observation is consistent with the definition of FDR = N_fp_/(N_tp_ + N_fp_) (**Supplementary Fig. 4**). When the modification of interest is highly abundant in a genome of interest, a relatively small number of false positives are associated with a low FDR as N_tp_ » N_fp_ (the native mPFC sample, **Fig. 1d**). However, when the modification of interest is of very low abundance in the genome (N_tp_ ∼ N_fp_ or N_tp_ « N_fp_), the false positive calls will result in a much higher FDR, meaning the vast majority of the called modifications are false positives (the mouse WGA sample in **Fig. 1d**). This raises caution for the use of nanopore sequencing to call 5mC from species with extremely low levels of 5mC. To avoid this pitfall, FDR analysis should be applied, instead of relying on arbitrary threshold on P_mod_, to reliably detect and interpret the called DNA modifications.

### Critical evaluation of the detection of 5mCpG, 5mCpH, 5hmC, and 6mA in mPFC

The above analysis was focused on 5mC across all sequence contexts in the mPFC sample. In mammalian genomes, 5mC is known to be mostly prevalent at CpG sites, but much less prevalent at non-CpG sites (i.e. 5mCpH, H=A/T/C): while certain brain cells have a high abundance of 5mCpH, most other cell types have very low levels of 5mCpH^28–30^. As expected, although 5mCpG calling has low FDRs in the mPFC sample, 5mCpH calling generally has much higher FDRs (**Fig. 1f**). This contrast between the reliability of 5mCpG and 5mCpH calling highlights the need for caution in the detection and interpretation of 5mCpH from various mammalian cell types. It also underscores the need to apply FDR evaluation specifically to modifications of interest within particular sequence contexts, rather than assuming equal reliability across all detected DNA modifications of the same type.

We further evaluated the use of nanopore sequencing for the direct detection of 5hmC, which is the oxidized derivative of 5mC and an intermediate in the DNA demethylation pathway. To achieve this, we applied the three-state model (dna_r10.4.1_e8.2_400bps_sup@v4.3.0, hereafter v4.3.0), which differentiate 5mC and 5hmC from unmethylated C, on the read-level modification analysis (**Methods**). For 5mC, FDR in the three-state model is notably lower than that of the two-state model. This suggests that the three-state model achieves higher 5mC prediction accuracy and enhances the differentiation between distinct modifications occurring on the same nucleotide (**Fig. 1d & g**). Compared to 5mC, 5hmC is much less abundant in mammal cells: while it is enriched in neurons, most other mammalian cells have very low levels of 5hmC^36,37^. As expected, a much lower 5hmC/C level was observed in mPFC than 5mC across all P_mod_ thresholds (**Supplementary Fig. 5**), consistent with previous estimations by LC-MS/MS^38^ (5mC/C: ∼4%-5% and 5hmC/C: ∼0.6%).

5hmC calling has a much higher FDR (**Fig. 1g-i, Supplementary Figs. 6 & 7**). Consistent results were observed in the mPFC sample with the DORADO model designed specifically for 5mCpG and 5hmCpG calling (“5mCG_5hmCG” field) (**Supplementary Fig. 8**). Notably, while 5hmCpG has a relatively low FDR, 5hmCpH calling showed the highest FDR, consistent with the previous studies using orthogonal enzymatic methods that show 5hmC enrichment in the brain is almost exclusively at CpG sites^31,36^ (**Fig. 1i, Supplementary Fig. 7**).

Next, we assessed the direct detection of 6mA, a DNA modification that is prevalent in bacteria but extremely rare in mammalian cells^12,14,15,20,21,39^. Although the official tools recently enabled 6mA calling, the FDRs estimated on the mPFC sample are ∼1 across P_mod_ thresholds, suggesting that the called 6mA events are essentially all false positives (**Supplementary Fig. 9)**. Amid the ongoing debate over 6mA in mammalian genomes^12,14,15^, this assessment highlights the critical need for the use of FDR to safeguard against unrecognized false positive calls in DNA modification detection that could mislead biological and biomedical studies, consuming years of research efforts and resources.

### Critical evaluation of the detection of 5mCpG, 5mCpH, 5hmC and 6mA in hLCL

Furthermore, we applied the FDR-based framework to a human lymphoblastoid cell line (hLCL, **Methods**). Read-level analysis on native nanopore reads showed 2.4%-4.9% 5mC/C level across the hLCL genome with the DORADO v4.3.0 model (**Supplementary Fig. 10**), consistent with previous characterization (∼2.5%-4.0%) as estimated by LC-MS/MS^40^. FDR evaluation showed a similar trend of decrease, from 0.76 to 0.08 as the P_mod_ threshold increased from 0.5 to 0.99 (**Fig. 1j**), suggesting 5mC calling is largely reliable as observed for native mPFC gDNA. However, FDR analysis of 5mCpH, 5hmC and 6mA calling showed that nanopore-sequencing-based calling of these three types of modifications are dominated by false positives in the hLCL sample, with high FDRs (∼1 across all P_mod_ thresholds for 5hmC and 6mA), consistent across versions of DORADO base calling models (**Fig. 1j-l, Supplementary Figs. 11-14**). These observations are highly consistent with previous studies that showed: (1) 5mCpH is rare in most mammalian cell types except certain brain cells^28–32;^ (2) the 5hmC/C level in hLCL is very low (only ∼0.001%) as reported by LC-MS/MS^40^; and (3) 6mA in hLCL is rare, consistent with our previous assessment of 6mA in hLCL using the independent PacBio sequencing^12,13^.

By comparing mPFC (moderate 5hmC enriched at CpG) and hLCL (very rare 5hmC), the FDR evaluation showed that 5hmCpG calls are largely reliable in mPFC, but not in hLCL (**Fig. 1i&1l**). These findings are consistent with previous estimates of 5hmC/C level by LC-MS/MS, which showed that hLCLs have much lower 5hmC/C levels compared to mPFC: ∼0.6% in mouse cerebral cortex^34,38^ vs. ∼0.001% in hLCL^40^. This comparison between different cell types highlights the distinct reliability of the same modification calling tool across samples with different levels of DNA modification.

### Benchmark of modifications detected by ONT data against independent enzyme-based methylome profiling methods in hPBMC and mPFC

To investigate the origins of false positives and characterize how modification detection rates vary across the genome, we compared ONT-derived calls against independent, enzyme-based methylome profiling methods. In addition to mPFC (enriched 5mC, moderate 5hmC), we evaluated a human peripheral blood mononuclear cell (hPBMC) line gDNA. Similar to hLCL, hPBMC gDNA exhibits a high FDR for 5hmC due to its low natural abundance ^41,42^, making it a great candidate for rigorous evaluation (**Fig. 2a**).

**Fig. 2.**
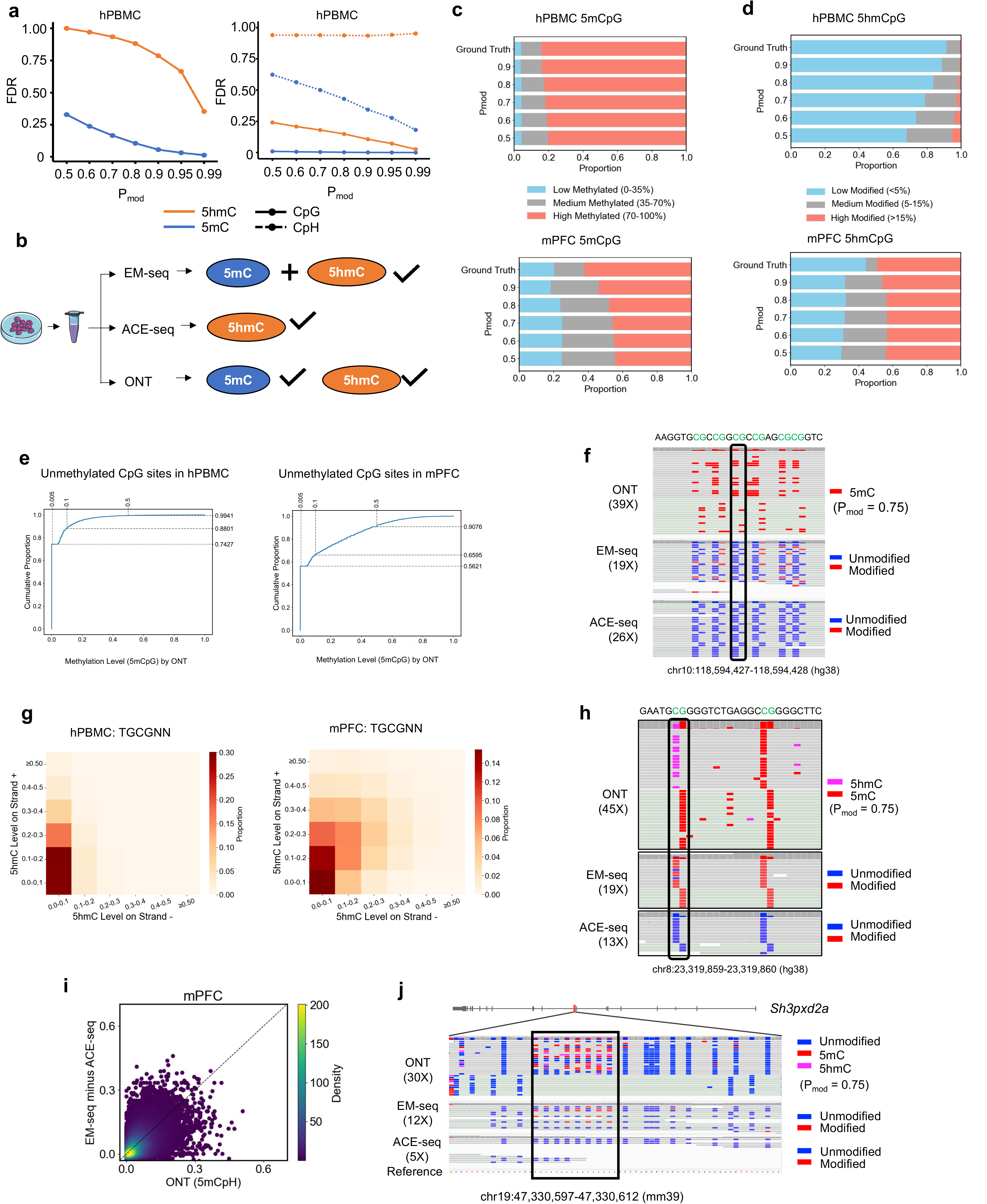
Benchmark of modifications detected by ONT data against independent enzyme-based methylome profiling methods in hPBMC and mPFC. a, FDR evaluation of modification calls made from the native hPBMC sample with the 5mC_5hmC model (v4.3.0). Left, 5mC and 5hmC calls. Right, 5mC and 5hmC calls made at CpG and CpH sites. b, Experimental design of benchmark pipeline. For hPBMC and mPFC gDNA, matched ONT data, EM-seq and ACE-seq were generated separately. EM-seq identifies but do not differentiate 5mC and 5hmC. ACE-seq specifically identifies 5hmC. ONT detects 5mC and 5hmC separately. c, Proportion of different 5mCG methylation levels in hPBMC and mPFC by ONT and EM-seq/ACE-seq. Three levels (Low, Medium and High) methylated regions (Ground Truth) were defined by enzyme-based methylome profiling methods (EM-seq and ACE-seq). Different P_mod_ threshold (From 0.5 to 0.9) were applied on ONT data analyses. d, Equivalent analysis for 5hmCG, performed as described for Fig. 2c. e, False-positive 5mCpG calls based on gold standard independent methods (EM-seq/ACE-seq). Cumulative proportion of CpG sites called by DORADO at loci with 0% methylation (unmethylated CpG sites) based on EM-seq/ACE-seq. Bins (0.005) show cumulative fractions; *x-axis*, ONT methylation levels; *y-axis*: cumulative proportion of loci below each threshold. f, Example of recurrent false-positive 5mCpG calls by ONT in the hPBMC sample at chr10:118,594,427-118,594,428 (hg38) CpG locus, not supported by EM-seq/ACE-seq, visualized in IGV. Regional sequencing coverage for ONT, EM-seq, and ACE-seq across the displayed IGV region is indicated in brackets. g, Proportion of 5hmCG on both strands at TGCGNN loci across hPBMC and mPFC genome from ONT data. *x-axis*, 5hmC level on strand -. *y-axis*, 5hmC level on strand +. False positive 5hmC sites were identified as CpG sites with 0% 5hmC levels in ACE-seq but ≥10% in ONT data. Equivalent analysis for all CG sites was shown in **Supplementary Fig. 18**. h, Example of repeatedly false positive ONT 5hmCG calls with a hemi-methylation pattern at chr8:23,319,859-23,319,860 (hg38) CpG locus in the hPBMC sample. Regional sequencing coverage for ONT, EM-seq, and ACE-seq across the displayed IGV region is indicated in brackets. i, Comparison of methylation level on (CA)n elements between EM-seq/ACE-seq and ONT in mPFC sample. Equivalent analysis for hPBMC was shown in **Supplementary Fig. 19**. j, Example of 5mCpH in (CA)n repeat loci at chr19:47,330,597-47,330,612 in mm39. Regional sequencing coverage for ONT, EM-seq, and ACE-seq across the displayed IGV region is indicated in brackets. Another example was shown in **Supplementary Fig. 20**.

We analyzed matched ONT data (with the DORADO v4.3.0 model), alongside two orthogonal enzymatic methods for each sample: enzymatic methyl-sequencing (EM-seq) ^43^, which detects the combined signal of 5mC and 5hmC, and APOBEC-coupled epigenetic sequencing (ACE-seq)^44^, which specifically isolates 5hmC (**Fig. 2b**). These enzymatic methods are widely considered the gold standards for 5mC and 5hmC detection.

As expected, we observed a general increase in concordance between ONT and enzyme-based methylome profiling as P_mod_ was increased from 0.5 to 0.9 for both 5mCpG and 5hmCpG across different modification levels (**Fig. 2c-d**). Regarding genomic context, the strongest concordance was found in high-abundance 5mCpG across both samples and moderate-abundance 5hmCG in mPFC (**Supplementary Fig. 15**). Conversely, low-abundance modifications were associated with a high rate of false positives in the ONT data. This was particularly pronounced for 5hmCpG in hPBMC, as well as for 5mCpH and 5hmCpH in both mPFC and hPBMC (**Supplementary Fig. 15**). Further analysis revealed that the Pearson correlation between ONT and EM-seq/ACE-seq measurements is driven primarily by CG motif density rather than read coverage (**Supplementary Fig. 15-17**).

Despite the general concordance, we identified specific loci where ONT repeatedly produced false-positive 5mCpG calls that were absent in EM-seq/ACE-seq data (**Figs. 2e-f**). Genomic context analysis revealed that 5hmC false positives frequently clustered at specific motifs (e.g., TGCGNN) in both hPBMC and mPFC samples. These calls exhibited a distinct strand bias, characterized by a 5mC call on one strand and a 5hmC call on the opposite strand (**Figs. 2g-h, Supplementary Fig 18**). As illustrated in **Figure 2h**, asymmetric 5hmC:5mC configurations at chr8:23,319,859-23,319,860 (hg38) CpG locus within the TGCGNN sequence context was observed across multiple ONT reads (45X), whereas EM-seq (19X) and ACE-seq (13X) identified the same site as symmetric 5mC. This false positive asymmetry is likely attributable to ambiguity in the machine learning–based modification caller, which can misclassify signals when the nanopore current associated with a given modification varies substantially across different sequence contexts ^4,45^. Regarding CpH sites, despite the elevated false-positive rates (compared to CpG sites), we identified a type of modification motif that are much more likely to be true positive, supported by both platforms. A notable example is the enrichment of 5mCpH within (CA)n repeats in the mPFC, a feature that is notably absent in hPBMC (**Figs. 2i–j, Supplementary Figs. 19-20**). This observation suggests a general strategy to enhance the reliability of modification calling by leveraging local detection density, based on the rationale that true positives are more likely to occur in clusters, whereas false positives arising from sequence-specific artifacts are less likely to co-occur across diverse neighboring sequence contexts.

### The use of more comprehensive negative controls to account for confounding DNA modifications

The above FDR analyses were all based on the use of WGA samples as negative controls. While WGA serves as very helpful negative control to capture technology-centric false positives, it does not consider the confounding effect among different forms of DNA modifications (**Fig. 1a)**. On this basis, we hypothesized that the abundant 5mC in CpG sites on the mammalian genome may further confound nanopore based detection of 5hmC which also locates in CpG context, leading to false positive 5hmC calls (5mC being called as 5hmC that are not represented by WGA samples). For example, the FDR of ∼0.4, as estimated using WGA as the negative control (**Fig. 3a**), appeared to support that 60% of 5hmCpG events called from hLCL are “reliable”. If this were true, it suggests 5hmC/CG levels of 0.047%-2.1%. Considering CpG dinucleotides account for 9.82% of total Cs (**Methods**), this equates to 0.0046%-0.21% 5hmC/C in the native hLCL genome (**Supplementary Fig. 14**), which is much higher than the 5hmC/C level (∼0.001%) estimated by LC-MS/MS and BS-seq/oxBS-seq technologies from the same cell type^40^. This alerted us to consider the abundant 5mCpG events in hLCL may have confounded the detection of 5hmC, leading to false positive 5hmCpG calls not captured by WGA.

**Fig. 3.**
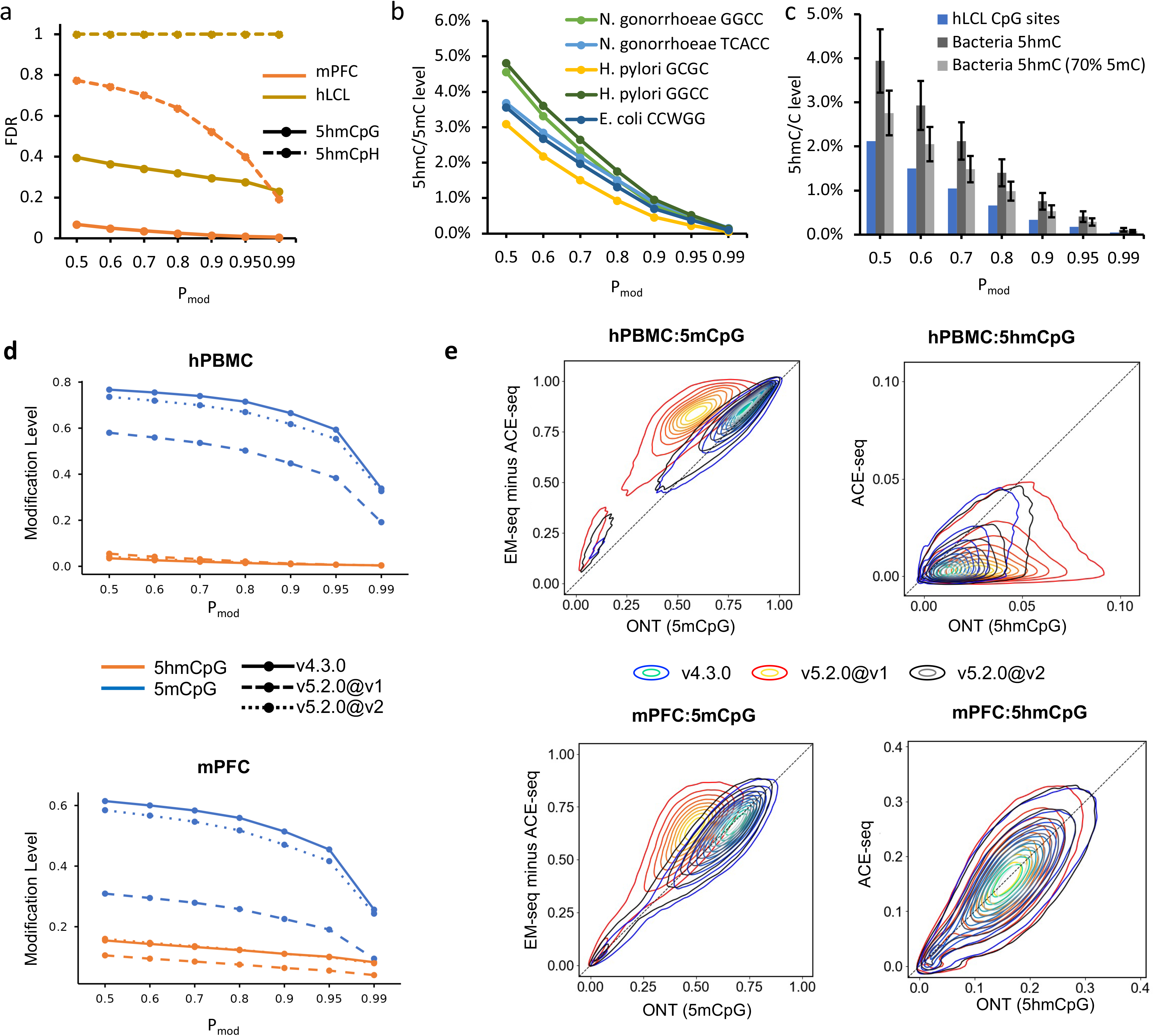
Comprehensive evaluations of confounding DNA modifications and different base call models. **a**, Comparison of 5hmC at CpG and CpH sites between mPFC and hLCL samples. **b**, False positive (FP) 5hmC/5mC calls from bacterial genomes with well-characterized 5mC motifs (nearly 100% methylated) yet free of 5hmC. **c**, The 5hmCG/CG levels detected in hLCL samples vs the FP 5hmC/5mC levels detected from bacterial 5mC motif sites that are free of 5hmC. The mean and standard deviation (represented by bars) of 5hmC levels were calculated from all 5mC motifs. FP 5hmC calls were estimated from 5hmC calls made from native bacterial reads alone, or in a mixture of 70% native bacterial reads and 30% WGA reads (**Methods**). d, Modification levels of 5mCpG and 5hmCpG among all CpG sites with different DORADO base calling models. e, Comparison of overlapping 5mCpG and 5hmCpG between ONT data analyzed with DORADO models and enzyme-based methods in hPBMC and mPFC by contour plots. x-axis, 5mCpG or 5hmCpG levels analyzed by DORADO model v5.2.0@v1, v5.2.0@v2 and v4.3.0. y-axis, 5mCpGs and 5hmCpG levels by enzyme-based methods. Pmod ≥ 0.75 was used for ONT-based analyses.

To test the hypothesis, we sequenced three bacterial strains with well-characterized 5mC motifs that cover all C contexts (CA, CT, CG and CC), but no 5hmC events (given the absence of associated TET enzymes^46^ or hydroxymethylases ^44^; see **Methods**). These bacterial data serve as helpful negative controls (abundant 5mC yet 5hmC free) that allow us to evaluate false positive 5hmC calls due to the confounding 5mC on the same context. Specifically, the *Neisseria gonorrhoeae* strain has GG5mCC and T5mCACC motifs^7,47^; the *Helicobacter pylori* strain has GG5mCC and G5mCGC motifs^45,47^; and the *Escherichia coli* K12 MG1655 strain has C5mCWGG (W=A/T)^47,48^. All of these motifs were nearly ∼100% 5mC methylated across the genome by their own methyltransferases^47,49–51^.

We assessed 5hmC calling in these bacterial data (**Methods**) and found that DORADO miscalls ∼3.94%-0.11% of 5mC as false positive 5hmC, when P_mod_ increases from 0.5 to 0.99 (with the v4.3.0 model). These false positive 5hmC/5mC levels in native bacteria are significantly higher than 5hmC/C levels in the matched WGA controls (**Fig. 3b, Supplementary Fig. 21**), supporting that these 5hmC calls were indeed false positives confounded by the abundant 5mC events. Importantly, the false positive 5hmC/5mC level miscalled from 5mC on bacterial genomes are higher than the 5hmCpG/CpG levels called from hLCL (**Fig. 3c, Supplementary Fig. 21**).

Even with a simulation of 70% 5mC/C levels among these motifs (to mimic the ∼70% 5mCpG/CpG level on human genome, see **Methods**), the rate of false positive 5hmC calls due to confounding with 5mC remains higher than 5hmCpG/CpG levels detected in the hLCL data (**Fig. 3c**). This result implies that although the WGA-based negative control estimated a seemingly low FDR for 5hmCpG calling from hLCL, the use of additional negative controls, high in confounding 5mC but 5hmC-free, provided critical estimation of high FDR (∼1, **Supplementary Fig. 22**), suggesting that the 5hmCpG events called from the hLCL sample are nearly all false positives. This analysis not only highlights that caution is needed when detecting 5hmC on a genome with abundant 5mC, but also the generally applicable need for more comprehensive negative controls in cases where different forms of DNA modifications can confound each other (**Fig. 1a**).

### Benchmark the latest ONT modification calling model using mPFC and multiple human cell lines and tissues

Besides the previously evaluated DORADO base calling model (v4.3.0), we also assessed two versions (v1 and v2) of a newer model (dna_r10.4.1_e8.2_400bps_sup@v5.2.0, hereafter v5.2.0@v1 and v5.2.0@v2) for modification calling, which is the latest release as of June 15, 2026 (**Supplementary Fig. 22-30**). We evaluated the detection of 5mC and 5hmC in both CpG and CpH contexts across mPFC, hLCL and several other human tissues using the modFDR framework (**Supplementary Figs. 23-24**).

We observed a consistent trend toward lower FDR in 5hmCpG with v5.2.0@v1 model whereas v5.2.0@v2 model exhibited FDR values between v5.2.0@v1 and v4.3.0 (**Figs 1h, 1k, Supplementary Fig 23-24**). For instance, in native mPFC gDNA data, increasing the P_mod_ threshold from 0.5 to 0.99 reduced the FDR for 5hmCG from 0.07 to 0.01 with the v4.3.0 model. In contrast, the v5.2.0@v1 model achieved a substantially tighter FDR range of 0.01 to 0.004, while v5.2.0@v2 yielded a range of 0.02 to 0.005 (**Supplementary Fig. 23-24**). Similar performance differences were observed in the native hLCL gDNA data, where the v4.3.0 model’s FDR decreased from 0.40 to 0.26 (**Fig. 1l**), while the v5.2.0@v1 model dramatically reduced the FDR to a range between 0.03 to 0.02 (**Supplementary Fig. 23**), and v5.2.0@v2 yielded an FDR range between 0.15 to 0.08 (**Supplementary Fig. 24**).

Existing literature indicates that brain tissues exhibit the highest 5hmC levels ^36–38^, while the kidney have lower (than brain) but higher levels compared to many other human tissues^52,53^. In contrast, hPBMC and hLCL typically show extremely low 5hmC levels^40^. While FDR evaluations with the v4.3.0 model effectively differentiate between these samples (**Supplementary Fig. 25)**, the v5.2.0@v1 model maintained a uniformly low FDR for 5hmCG regardless of the known biological levels (**Supplementary Fig. 26)**, indicating that v5.2.0@v1 model may exhibit lower robustness in detecting 5hmCpG modifications. For a better understanding and comprehensive evaluation, we utilized the gold standard enzyme-based EM-seq and ACE-seq on mPFC and hPBMC and compared them with the ONT data generated by the updated v5.2.0 models.

We first checked the false positives for 5mC and 5hmC previously detected with the v4.3.0 model. False positive 5mCG and 5hmCG detections persisted in v5.2.0 models on both mPFC and hPBMC gDNA at loci confirmed to have 0% modification by EM-seq/ACE-seq (**Supplementary Fig. 27-30**). Meanwhile, false positive 5hmC signals associated with asymmetric 5hmC:5mC configurations at TGCGNN sites remain in both samples (**Supplementary Fig. 29-30**). False-positive 5hmCG calls that were associated with confounding high levels of 5mCG also persisted. Specifically, ONT data showed increased 5hmCG predictions as 5mCG levels rose, despite these sites being identified as pure 5mCGs by gold-standard EM-seq/ACE-seq (**Supplementary Fig. 31**).

We next evaluated the concordance of 5mC/5hmC within CpG and CpH contexts between ONT data (using v4.3.0, v5.2.0@v1 and v5.2.0@v2) and the enzyme-based profiling. For 5mCpG sites, we observed a significance decrease in the global 5mCpG/CpG level across mPFC, hPBMC and other human samples with the v5.2.0@v1 model (**Fig. 3d, supplementary Fig. 32**), whereas v5.2.0@v2 maintained levels comparable to v4.3.0. For instance, using a P_mod_ 0.8 in hPBMC, the estimated 5mCpG/CpG ratio dropped from 0.72 with v4.3.0 to 0.50 with v5.2.0@v1, compared to 0.67 with v5.2.0@v2. Compared to the gold-standard EM-seq/ACE-seq, v4.3.0 and v5.2.0@v2 model showed strong concordance with gold-standard measurements, whereas v5.2.0@v1 exhibited high deviation in both mPFC and hPBMC (**Fig. 3e**). These findings indicate a notable increase in false-negative 5mCG calls (true 5mC sites not detected) in the v5.2.0@v1, an issue that was largely corrected in v5.2.0@v2.

For 5hmCG, although mPFC gDNA showed high concordance with enzyme-based profiling across all ONT models (v4.3.0, v5.2.0@v1 and v5.2.0@v2), the low-abundance 5hmCG in hPBMC ONT data exhibited greater discrepancy (false positives), particularly with the v5.2.0@v1 model (**Fig. 3e**). Furthermore, compared with the v4.3.0, the v5.2.0@v1 model predicted a slight increase in 5hmCG/CpG for kidney but a decrease for brain (**Supplementary Fig. 32**). Coupled with the reduced 5hmCpG prediction in human WGA sample (**Supplementary Fig. 32**), this difference between v5.2.0@v1 and v4.3.0 partially explains to the uniformly low FDR for 5hmCpG in modFDR evaluation of brain and kidney tissues. Collectively, these analyses against benchmark EM-seq/ACE-seq data suggest that the v5.2.0@v1 model may possess lower robustness in detecting 5hmCpG modifications across different tissue types.

Regarding the CpH context, v4.3.0, v5.2.0@v1 and v5.2.0@v2 models showed discordance for 5mCpH and 5hmCpH detection compared to the gold-standard EM-seq/ACE-seq. While the model v4.3.0 tended to over-estimate (false positives), the v5.2.0@v1 model appeared more prone to underestimating CpH modifications (false negatives). Although v5.2.0@v2 yielded intermediate estimates, it still produced high false positive especially 5hmCpH in mPFC. These findings demonstrate that accurate modification calling in CpH contexts remains a key challenge for Dorado models. (**Figs 1h, 1k, Supplementary Figs 23 & 24**).

In summary, while Dorado v5.2.0@v1 introduced substantial structural changes compared to v4.3.0, the revised v5.2.0@v2 model offers an intermediate performance profile that resolves some, but not all limitations. Across all tested models, several critical challenges remain: (1) systematic false positives in both 5mCG and 5hmCG persist, including strand-biased 5hmC calls at specific motifs such as TGCGNN; (2) false-positive 5hmCG calls can be confounded by high 5mCG abundance; (3) the v5.2.0@v1 is showing high discordance with EM-seq/ACE-seq measurements, specifically an elevated false-negative rate for 5mCpG, but this systematic bias was largely corrected in v5.2.0@v2; (4) all models are exhibiting lower robustness in detecting 5hmCpG modifications; and (5) reliable detection of 5mC and 5hmC in the CpH context remains a challenge.

## Discussion

In this study, we have critically assessed the reliability of nanopore sequencing for detecting multiple forms of DNA modifications, demonstrating that false positives can significantly influence data interpretation, especially for low-abundant modifications. This raises the need for caution in the current common practice where arbitrary thresholds are used for calling DNA modifications from nanopore data, which can confound data interpretation and mislead downstream biological analysis.

To address these challenges and guide both users and method developers, we have presented modFDR, a framework that emphasizes the use of rationally designed negative controls (capturing both general background and confounding modifications) and FDR analysis (**Fig. 1a**). Using modFDR, we demonstrated that the abundance of modification plays a critical role in the FDR evaluation. Specifically, we applied this framework to assess the calling of multiple modification types (5mC, 5hmC, and 6mA) at different genomic contexts (CpG and CpH), and we made observations consistent across both mouse and human samples using various versions of DORADO base calling models (v4.2.0, v4.3.0, v5.0.0 and latest v5.2.0) as of June 15, 2026.

Traditional whole-genome bisulfite sequencing (WGBS) methods are often influenced by various artifacts, including experimental issues (e.g., insufficient or prolonged bisulfite conversion, PCR amplification) and computational challenges inherent in next-generation sequencing (e.g., sequencing bias, alignment bias) ^54^. Nanopore sequencing, being amplification-free, has been proposed as a superior method for modification detection ^54^. In our study, we utilized the latest R10.4.1 flowcell for nanopore sequencing. The R10 chemistry was specifically designed to deliver the highest consensus accuracy and has been reported to show the highest genomic alignment rates and correlation with WGBS when compared to the older ONT R9 flowcell and the PacBio platform ^55^. However, our investigation demonstrated that, even with the highly accurate R10 sequencing chemistry and the latest base-calling models, this amplification-free method still needs improvement in reducing false positive detection. Therefore, modification calls must be reliably quantified using modFDR analyses before biological interpretation. For further validation, integrating nanopore data with alternative technologies, such as PacBio SMRT sequencing or NGS platforms coupled with modification-sensitive restriction enzymes, remains a more robust strategy for definitive confirmation.

The broad utility of modFDR is particularly helpful because DNA modification levels are known to exhibit significant biological variation across different tissues, cell types, and developmental stages, as observed in our study (**Supplementary Fig. 3**) and previous research ^34,38,56^.This biological variability highlights the crucial importance of performing modFDR analysis specifically for the sample of interest, rather than assuming reliable detection efficiency or modification abundance levels based solely on prior knowledge.

For 5mC, while detection at CpG sites in the mammalian genomes is generally reliable with a low FDR, the same cannot be said for CpH sites, because 5mCpH is typically rare in most mammalian cells^28–32^. Our modFDR analyses are consistent with previous studies that demonstrated lower error rate for CpG compared to CpH detection in both vertebrates ^57^ and plant^58^ based on low-pass nanopore sequencing (Skim-seq) for measurement of global methylation levels, further validating 5mC detection in CpH contexts is a recurring challenge across different biological systems. Whereas Skim-seq is intrinsically limited to aggregate estimations of global modification levels, the modFDR framework offers broad utility for locus-specific analysis by leveraging a true negative control baseline.

For 5hmC, although high reliability can be achieved in cell types where they are relatively abundant, such as the mPFC, human brain and kidney tissues, high false positive rates are expected in most other mammalian cell types where they are rare. For 6mA, which is prevalent in bacteria but largely absent in mammalian genomes^12,14,15^, our FDR analysis showed that it cannot be reliably detected yet using nanopore sequencing even with the latest official model release. The generally high FDRs associated with 5mCpH, 5hmC, and 6mA calling from most mammalian cell types are in great contrast with the high detection accuracy (97.9% and 97.5% for 5mC/5hmC and 6mA all context detection, respectively) for DNA modifications reported by the nanopore sequencing official tool DORADO (https://nanoporetech.com/platform/accuracy/#variant-calling). This is essentially because the DORADO models were all developed and evaluated using training datasets with predefined ratios of modified and non-modified bases, which do not reflect the true biological variability in modification abundance in mammalian cell types.

Previous studies reported that 6mA in metazoa may originate mostly from bacterial contamination (which can significantly skew LC-MS/MS ^12,14,15,20,21^. Such massive contamination often leads to incorrect assumptions about the modification abundance. For example, although commensal/environmental bacteria introduced substantial contamination in 6mA studies in mammalian genomes, an equally fundamental challenge was the actual absence or ultra-low levels of 6mA in mammalian genomes^4,12^. In this context (low 6mA abundance in mammalian genomes), the high FDR of nanopore sequencing for detecting 6mA, combined with the use of arbitrary cut-offs, led to numerous 6mA calls (from the mammalian genome) that appeared “consistent” with LC-MS/MS estimates. If previous studies that overestimated 6mA had used modFDR analysis with negative controls, they would have recognized the unreliable 6mA calls from the mammalian genome, in stark contrast to the LC-MS/MS estimates, even without knowledge of massive contamination.

Our work highlights the urgent need for the community to incorporate modFDR framework, along with rationally designed negative controls in their nanopore-based DNA modification mapping studies, especially for method development and evaluation. It is worth noting, while our study emphasizes the use of FDR to quantify false positives in practical applications, we fully recognize the value of established metrics, such as AUC (Area Under the Curve), FPR, FNR (False Negative Rate), sensitivity/recall. However, these traditional metrics may not be sufficient for real-world applications involving low-abundance modifications, specific genomic context, or species with confounding modifications. Indeed, our comparison of between the newer and older DORADO models highlights the importance and complementarity between using both FDR and false negative rates. To ensure rigorous method development and evaluation, FDR should be included as a crucial metric, especially within the context of physiologically relevant levels of DNA and RNA modifications (**Fig. 1a, supplementary Fig. 4**).

For biological studies and biomedical applications, our study supports the general reliability of 5mCpG mapping across mammalian cell lines, but limited applicability of existing 5hmCpG mapping methods to brain tissues or other tissues with high 5hmC abundance. In addition, our findings indicate a critical need for further efforts to incorporate data with physiologically relevant levels of modifications in the model training and evaluation, along with matched data generated using gold-standard methods such as EM-seq ^43^ and ACE-seq ^44^. Before new reliable methods are developed and critically assessed for mapping low-abundance modifications, nanopore sequencing should be prioritized for mapping abundant, rather than rare, modifications in biological studies and biomedical applications.

We demonstrated performance difference across different versions of the updated DORADO model (v5.2.0). While the model v5.2.0@v1 exhibited a notable rise in false negative 5mCpG calls, v5.2.0@v2 largely resolved this systematic bias, yielding performance comparable to the older v4.3.0 (**Fig. 3e**; **Supplementary Fig 33**). Despite these updates, challenges persist across all model versions, including prevalent systematic false positives for 5mCG and 5hmCG in mammalian cells (**Supplementary Figs. 27–31**), and unreliable modification calling within the CpH context (**Supplementary Fig. 33**). These findings suggest that, despite recent model releases, substantial algorithmic advancements are still required before the model can function as a reliable tool across diverse modification types and genomic contexts.

On the other hand, we observed that applying the 5mC_5hmC model field in v5.2.0@v1 to bacterial data yielded abnormally low 5mC levels (ranging from 30% to 80% at P_mod_=0.5) within established 5mC motifs. Given that these motifs are nearly 100% methylated by endogenous methyltransferases ^47,49–51^, this suggests a significant bias when applying the 5mC_5hmC field to bacterial genomes **(Supplementary Fig. 34)**. Conversely, the 4mC_5mC model field yielded 5mC calls for 80%–90% of cytosines (P_mod_=0.5) within the same motifs, indicating a relatively more accurate estimation of 5mC within a bacterial context **(Supplementary Fig. 34)**. The 5mC_5hmC and 4mC_5mC fields demonstrate distinct advantages in mammalian and bacterial contexts, respectively, which suggest these models were likely trained on different datasets. Therefore, they must be selected and applied with caution based on the specific biological applications and host genomes.

Furthermore, we emphasized the importance of employing comprehensive negative controls that account for the confounding effects of abundant modifications on the detection of rarer ones, by demonstrating a significant level of false positive 5hmC called from abundant 5mCpG methylation in human cells (**Fig. 3b & c**). This result is consistent with findings by Halliwell et al., who detected high levels of false positive 5hmC sites from nanopore data originating from enzymatically methylated, 5mC-positive human gDNA^59^, at rates far exceeding the natural 5hmC levels previously estimated for that human cell line^60^. Such confounding problems also exist in the diverse RNA modifications, e.g. ambiguous predictions between m6A and m1A, which occur on the same nucleotide ^61^.

Step-by-step methodological guidelines for modFDR evaluation of modification types are provided in **Supplementary Fig. 35.** As the nanopore sequencing community aims to advance toward the simultaneous detection of various DNA modifications, such as DNA methylation, DNA damage, and abasic sites^62,63^, the potential for cross-confounding effects cannot be overlooked. This caution also extends to the direct nanopore sequencing of RNA, where the diverse landscape of RNA modifications presents similar challenges^4,64,65^. A recent systematic evaluation of 86 computational tools for RNA modification identification revealed that factors such as modification levels and sequence motifs significantly confound the accurate detection of m6A, m5C and other RNA modifications^61^. Consequently, no single tool currently excels across all key performance metrics, including accuracy, biological validity, robustness, and computational efficiency ^61^. Given the practical complexities encountered in real-world applications, modFDR could serve as a valuable complementary tool for the empirical evaluation of computational methods, particularly when applied to specific samples in RNA modification research. Especially, additional negative controls are required to capture the interference between different RNA modifications (as we have demonstrated using 5mC and 5hmC on DNA in this study). This is even more critical for the mapping of RNA modifications, given its greater diversity.

Although our study focused on nanopore, modFDR can also be helpful modification analyses based on other technologies, such as PacBio SMRT sequencing, which has been broadly used in other modification analysis such as 6mA. While our manuscript was under revision, PacBio (April 2025) announced 5hmC detection capabilities based on a recent study^66^ and promoted broad adoption by the user community. In this context, modFDR, will not only help nanopore sequencing users but also PacBio sequencing users, making an even broader impact.

It is worth noting that modFDR is not only applicable to native whole-genome sequencing but also to sequencing DNA samples following modification enrichment. For example, although 5hmC is low in most mammalian cells, a future strategy to overcome false positives is to combine antibody-based enrichment—well established in the epigenetics and epigenomics communities—with nanopore or PacBio sequencing. The elevated levels of 5hmC post-enrichment are expected to reduce the proportion of false positive calls, which can be rigorously quantified using the modFDR framework.

In conclusion, the implications of our findings are particularly significant given the widespread application of nanopore sequencing in fields ranging from fundamental biological research to clinical diagnostics, including the analysis of cell-free DNA and RNA in liquid biopsies^67,68^. Detecting low-abundance modifications remains a considerable challenge, underscoring the need for a rigorous, transparent framework to evaluate the reliability of detected events.

## Acknowledgments

We thank Magdalena Ksiezarek, Yujie Liu and Yangmei Li for their help with nanopore sequencing library preparation and bacterial genomic DNA samples that we used as negative controls in this study. This work was supported by the Biorepository and Pathology Core at the Icahn School of Medicine at Mount Sinai; we thank Dr. Rachel Brody, the core staff, and the biorepository participants. We thank Kenny Jing Lo, Drs. Indu Saini and Kennedy Okhawere, Jewel Bamby, and Chitra Hindnavis from Dr. Ketan Badani’s laboratory for their assistance with kidney sample and data collection. We also thank the Neuropathology Brain Bank & Research CoRE—specifically Drs. Jamie Walker and Claudia De Sanctis, and Emma Thorn—for providing brain samples and collection support.

This work was supported by grants no. R35 GM139655 (G.F.), R01 HG011095 (G.F.), and R01 HG010646 (R.M.K. and G.F.) from the National Institutes of Health. C.E.L. was supported by F31 HG012892. This work was also supported by a Development Award as part of the Mount Sinai ADRC (NIH grant P30 AG066514). G.F. is a Hirschl Research Scholar by Irma T. Hirschl/Monique Weill-Caulier Trust and a Nash Family Research Scholar. This work was supported in part by the staff and resources of Department of Scientific Computing at the Icahn School of Medicine at Mount Sinai. Y.K. was an employee at Icahn School of Medicine at Mount Sinai. During this project, she assumed a new role as a Professor at the Advanced Institute for Life and Health, Department of Medicine, Southeast University (SEU) in Nanjing, China, where she was supported by her start-up fund (RF1028623368), the Fundamental Research Funds for the Central university (2242025F10004) and Southeast University Interdisciplinary Research Program for Young Scholars (2024FGC1004) from SEU.

## Data availability

Sequencing data generated in this study (**Supplementary Table 1**) have been deposited to Sequence Read Archive (SRA) data under BioProject ID PRJNA1177994.

## Author contributions

Study design: Y. K and G.F.; Data analysis: Y.K. and H.C.; Data processing: Y.K., Y. Z. and H.C.; Cell culture and sequencing: E.A.M.; Data interpretation: Y. K., Y. Z., H. C., E.A.M., C.E.L., Y. F., M. N., X-S Z., R. M. K. and G. F.; Supervised research: G.F.; Wrote first draft of paper: Y.K. and G.F.; Approved paper: all authors.

## Materials and Methods

### Mouse Prefrontal Cortex (mPFC)

10-12 mg PFC tissue was extracted from healthy female C57BL6 mice (Jackson Laboratory, ME, USA), aged 14-16 weeks old. Mice were reared in a specific pathogen-free (SPF) vivarium at Rutgers University’s School of Public Health animal facility by methods previously described^69^. Each brain was quickly removed and placed in an ice-cold petri dish, and prefrontal cortex (PFC) tissue was carefully extracted by brain dissection. The mPFC samples were flash-frozen individually in dry ice and stored at -80°C for subsequent DNA extraction. DNA isolation was carried out by a Wizard Genomic DNA Purification Kit (Promega Corporation, WI, USA), following the kit recommendations, including the optional RNase treatment. Another mPFC sample (mPFC2) was collected from a different mouse individual using the same protocol.

### Human Lymphoblastoid Cell Lines (hLCL)

Human lymphoblastoid cells samples were originally obtained from Coriell Institute for Medical Research (Coriell; cat# GM 12878, NJ, USA) and the methods utilized for LCL samples were as detailed previously^7^.

### Human Peripheral Blood Mononuclear Cell (hPBMC)

hPBMC (SER-PBMC-200 from ZenBio, Durham, NC, USA; Cell count >200x10^6^ PBMCs; Lot# PBMC012721B) was used for gDNA isolation. The sample came on ice as a 15mL conical of cells in serum, and cells were collected by spinning at 350xg/7’/4C. The supernatant was aspirated, and the pellet was flash-frozen. DNA isolation of the hPBMC sample was conducted using a Qiagen DNeasy Blood and Tissue kit (Qiagen #69504 Germantown, MD, USA) by manufacturer’s protocol, with the optional RNase A treatment. Samples were eluted in AE buffer.

### Human brain and kidney samples

Two postmortem frozen brain samples (NPBB251 and NPBB275) from temporal lobe adjacent to hippocampus were obtained from the Neuropathology Brain Bank & Research CoRE at MSH (RRID: SCR_027565). Another postmortem brain sample (FE1) is from Brodmann Area 4 (BA4) region, part of the pre-central gyrus. Two kidney tissues (kc_48034 and kc_48305) were obtained from the Biorepository and Pathology CoRE from surgeries conducted by the Department of Urology at MSH (IRB #19-00152). Samples were processed by the biorepository and provided flash frozen.

Genomic DNA extraction from kidney and brain tissues was performed using the Wizard Genomic DNA Purification Kit (Promega, Madison, WI, USA) following the manufacturer’s animal tissue protocol (sub-procedure 3D). Input material ranged from 50-250 mg for kidney samples and up to 500 mg for brain tissue. The protocol included the optional RNase digestion step. For larger tissue amounts, reagent volumes were proportionally increased at each step to maximize DNA yield. Extracted DNA was eluted in 0.1× IDTE buffer (IDT, Coralville, IA, USA), prepared by diluting the stock TE solution 1:10 in nuclease-free ultrapure water (ThermoFisher Scientific, Waltham, MA, USA).

#### Neisseria gonorrhoeae FA 1090

*N. gonorrhoeae* FA1090 was obtained from American Type Culture Collection (ATCC, cat# 700825, VA, USA) and the methods utilized for *N. gonorrhoeae* were detailed as previously described^7^.

#### Escherichia coli K-12 MG1655

*E. coli* K-12 MG1655 sample and methods were detailed using methods detailed previously^7^.

#### Helicobacter pylori JP26

*H. pylori* strain JP26 was originally isolated from a gastric cancer patient in Japan^70^, maintained as detailed previously^71^, and stored at -80°C. To obtain a working stock, the *H. pylori* strain was grown at 37°C in 5% CO_2_ on Trypticase soy agar (TSA) plates with 5% sheep blood (ThermoFisher Scientific # R01200, MA, USA) for 3 days, then some of the bacterial lawn was transferred to a new plate and incubated at 37°C in 5% CO_2_ for 2 days. Finally, bacteria were collected in a 1ml PBS rinse of the plate for DNA extraction with the Qiagen DNeasy Blood and Tissue Kit (Qiagen# 69504, MD, USA).

### Whole genome amplification (WGA)

REPLI-g multiple displacement amplification was carried out for *E.coli*, hLCL, mPFC, mPFC2 gDNA samples by the methodology described previously^72^. For the *E. coli,* mPFC2 samples, an additional sample was created for each (designated as 2-round WGA samples) by subjecting the first round of WGA samples to a second round of REPLI-g amplification. The rationale was to compare the two rounds of WGA to ensure DNA amplification was adequate and the WGA samples serve as reliable negative controls.

### ONT sequencing

Sequencing was primarily performed with PromethION P2 solo and MinION Mk1b and Mk1c instruments, R10.4.1 flow cells from Oxford Nanopore Technologies as described previously^72^; Ampure bead incubations were extended to 45 minutes improve elution. The rapid kit generated mPFC2 library was sequenced on a MinION Flow Cell (R10.4.1) until the estimated data yield reached 5GB.

### ACE-seq and EM-Seq library preparation and sequencing

For each sample undergoing ACE-seq, 20 ng of either purified mouse or human gDNA was mixed with 1% of unmethylated λ gDNA spike-in and 1% 5mCpG methylated plasmid spike-in and brought to a total volume of 50 µL in low-TE buffer. This input DNA was then sheared to 350 bp using a Covaris M220 ultrasonicator, end-repaired, and ligated to modified Y-shaped adaptors (forward: ACACTCTTTCCCTACACGACGCTCTTCCGATC*T, *indicates phosphorothioate, reverse: GATCGGAAGAGCACACGTCTGAACTCCAGTCA, all Cs are 5pyC, IDT) using the NEBNext UltraII DNA Library Prep Kit (NEB, E7645) according to the manufacturer’s instructions. The sample was then purified using a 1.2x left-sided SPRI cleanup (Beckman Coulter, B23317) according to the manufacturer’s instructions and eluted in 17.6 µL of nuclease-free H_2_O (nfH_2_O). Next, 16.6 µL of the eluent was treated with 1 µL bGT (NEB, M0357) in a reaction containing 0.4 µL UDP-glucose (NEB, S2200) and 2 µL of 10x CutSmart Buffer (NEB), and incubated at 37 °C for 1.5 h. Following conversion, samples were purified using a 1.2x left-sided SPRI cleanup and eluted in 17 µL of nfH_2_O. Subsequently, 16 µL of eluent was snap-cooled by adding 4 µL of formamide (Thermo Fisher, 17899), heating to 85 °C for 10 min, and immediately placing the sample on ice. The resulting 20 µL of snap-cooled mixture was combined with 80 µL of a master mix containing 68 µL of nfH_2_O, 10 µL of APOBEC 10x buffer (NEB, E7134), 1 µL of BSA (NEB, E7135), and 1 µL of APOBEC (NEB, E7133), and incubated at 37 °C for 3 h. The deaminated mixture was then purified by 1.2x left-sided SPRI cleanup and eluted in 23.5 µL of nfH_2_O. Next, 22.5 µL of eluent was combined with 25 µL of 2x KAPA HiFi HotStart U^+^ ReadyMix (Roche, 50–196–5287) and 2.5 µL of NEBNext Multiplex Oligos for Enzymatic Methyl-seq (NEB, E1720), and amplified using 8 PCR cycles (PCR program: initial 30 seconds at 98 °C; followed by repeat cycles of 10 seconds at 98 °C for denaturation, 30 seconds at 60 °C for annealing, 60 seconds at 65 °C for extension). Amplified libraries were purified using a 0.8x left-sided SPRI cleanup and eluted in 11 µL nfH_2_O to yield final libraries. Final libraries were quantified using a Qubit HS kit (Invitrogen, Q32851), assessed for quality using a High Sensitivity D1000 ScreenTape (Agilent, NC1786959), and pre-sequenced on an Illumina MiSeq to assess spike-in conversion efficiency.

For each sample undergoing EM-seq, libraries were prepared using the NEBNext Enzymatic Methyl-Seq kit (NEB, E7120L) according to the manufacturer’s instructions. As with ACE-Seq, 20 ng of either purified mouse or human gDNA was mixed with 1% of unmethylated λ gDNA spike-in and 1% 5mCpG methylated plasmid spike-in and brought to a total volume of 50 µL in low-TE buffer. The DNA mixture was then sheared to 350 bp using a Covaris M220 ultrasonicator and used as input for EM-Seq library preparation. Following end-repair, adaptor ligation, and enzymatic conversion, libraries were amplified using 8 cycles of PCR with KAPA HiFi HotStart U^+^ ReadyMix (Roche, 50–196–5287) mix, as above, and purified using a 0.8x left-sided SPRI cleanup, with elution in 11 µL nfH_2_O to yield final libraries. Final libraries were quantified using a Qubit HS kit (Invitrogen, Q32851), assessed for quality using a High Sensitivity D1000 ScreenTape (Agilent, NC1786959), and pre-sequenced on an Illumina MiSeq to assess spike-in conversion efficiency.

### Read-level modification analysis for mPFC and hLCL

Modified base calling was performed using DORADO (v0.5.3, v0.7, and v1.0.2; https://github.com/nanoporetech/dorado) against the genome reference for the (mm39) and human (hg38). Modification models were selected based on the specific modification of interest and the analytical goals (**Supplementary Table 2**). Counts of modified and unmodified bases were estimated using modkit (v0.2.6 and v0.3.1; https://nanoporetech.github.io/modkit) to manipulate modBAM generated by DORADO. To evaluate the FDR across different P_mod_ thresholds, we used the *modkit summary* function to calculate the candidates (%) that passed each specific threshold, for both native and WGA samples, separately. Specifically, modkit v0.2.6 was used for modBAM files produced by DORADO v0.5.3 and v0.7 with base calling models prior to v5.0.0. modkit v0.3.1 was used for modBAM files generated by DORADO v1.0.2 with the v5.2.0@v1, as well as Dorado v1.4 with the v5.2.0@v2 base calling models (**Supplementary Table 2**). All downstream modification analyses are restricted to reads that mapped to the respective reference genomes.

For the both mPFC and hLCL datasets, genome wide modification analyses were conducted using modkit summary with *-f 0.001*, and modification probability thresholds (P_mod_) were adjusted from 0.5 to 0.99 via the *--filter-threshold* parameter. For mPFC2 and the DORADO (v0.7)-processed subsamples of mPFC and hLCL (**Supplementary Table 1**), modified and unmodified base counts were estimated using *modkit summary.* These analyses were performed with the *--no-sampling* flag and adjusted P_mod_ ranging from 0.5 to 0.99 applied via the *--filter-threshold* parameter. Only sites that successfully passed the P_mod_ threshold (either confident unmodified or confident modified) were included for further calculation. To ensure comparability of modification levels across different thresholds, we first calculate the total count of all modification codes and canonical bases, by summing up *all_counts* output by modkit, denoted as S_all_. For each modification type of interest, such as 5mC, we then determine the number of modified calls passing a specific threshold (*pass_count* by modkit), referred to as S_pass_. The modification level is subsequently calculated by dividing S_pass_ by S_all_. This approach normalizes the modification levels, enabling accurate and consistent comparisons across various thresholds.

### False discovery rate (FDR) calculation

The FDR corresponding for a specific P_mod_ threshold was estimated by comparing the global distribution of modification calls obtained between native DNA and WGA (methylation-free) samples. *modkit summary* function was used to generate statistics for modified and unmodified base counts. The fraction of candidate sites passing a given P_mod_ threshold was calculated independently for native and WGA samples. Specifically, for a given threshold (thres), the FDR was defined as:

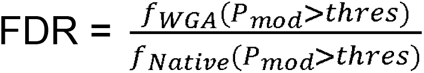

where P_mod_ denotes the classification probability by DORADO software for a specific base. f_WGA_(P_mod_>thres) is the fraction of sites called as methylated in the WGA sample out of total calls at that threshold. f_native_(P_mod_>thres) is the fraction of sites called as methylated in the native sample out of total calls at that threshold. Unmethylated sites with a P_mod_ value below the threshold in either sample were excluded from the FDR estimation. We evaluated P_mod_ thresholds ranging from 0.5 to 0.99 and capped the FDR at 1.0 to maintain conceptual consistency.

### Simulation of 5mC level of different magnitude orders on mPFC sample

To create 5mC levels of different magnitude orders, native mPFC and mouse WGA samples were randomly subsampled and mixed to create 5mC levels of different magnitude orders. Briefly, reads were randomly subsampled from two samples using samtools^73^ (v1.17) to create mock samples with 10M reads in total for each. Three independent simulations were performed to exclude stochastic noise during random subsampling. To mitigate coverage bias between native mouse and WGA samples, only regions with overlapping 5–30× coverage between two samples were used for subsampling simulations. Read counts from two samples were normalized based on the average read lengths to ensure the ratio of yields within the mock samples. Fraction of modification calls were performed using *modkit summary -f 0.01* with adjusted P_mod_ (from 0.5 to 0.99) applied on *--filter-threshold*.

### Genome-wide CpG and CpH site identification on mPFC and human tissue samples

Genome-wide counts of CpG dinucleotides and total cytosines were calculated using seqtk (v1.3-r106). CpG and CpH sites were identified using *modkit motif-bed reference.fasta CG/CH 0*. We used *modkit --include-bed* to restrict analysis to base modification probabilities overlapping these genomic intervals (CpG or CpH sites). The CpG and CpH modification levels and FDR analysis were performed as the methods described above.

### Data processing and methylation calling for EM-seq and ACE-seq

Raw EM-seq and ACE-seq reads were trimmed to remove low-quality bases and adapter sequences using Trim Galore (v0.6.6; https://github.com/FelixKrueger/TrimGalore), and data quality was assessed with FastQC (v0.11.9; http://www.bioinformatics.babraham.ac.uk/projects/fastqc). Reference genome indexes were generated using Bowtie2 (v2.4.4)^74^. The trimmed reads were aligned to the reference genomes using Bismark (v0.22.3)^75^, and PCR duplicates were subsequently removed with the Bismark deduplication module (deduplicate_bismark). Methylation level signal at each site was calculated as % of C/(C+T).

### Pairwise comparisons between EM-seq/ACE-seq and ONT

Genomic sites shared among ONT, EM-seq, and ACE-seq datasets were used for pairwise comparisons. The genome-wide ONT data depth for mPFC and hPBMC is 32.7X and 33.6X respectively. For comparison with ONT 5mC calls, the methylation signals were defined as the EM-seq signals minus ACE-seq signals, whereas ACE-seq signals were used directly for comparison with ONT 5hmC calls. For regional methylation level comparisons across the genome, methylation levels were aggregated within non-overlapping 5-kb genomic bins.

Because the hPBMC and mPFC EM-seq and ACE-seq datasets differed in sequencing depth (hPBMC EM-seq, 31.2×; hPBMC ACE-seq, 39.0×; mPFC EM-seq, 9.0×; mPFC ACE-seq, 8.0×), different coverage thresholds were applied. For hPBMC, 5-kb regions with ≥5× CpG/CpH coverage shared among ONT, EM-seq, and ACE-seq were used for pairwise comparisons. In mPFC, substantially fewer 5-kb regions had ≥5× CpG/CpH coverage in both the EM-seq and ACE-seq datasets. To increase the number of genomic regions available for whole-genome comparison, we performed a downsampling analysis of high-coverage mPFC EM-seq and ACE-seq regions and confirmed that regional methylation levels at 1× CpG/CpH coverage remained highly correlated with those obtained at the original higher coverage (Supplementary Fig. 36). Based on this analysis, regions with ≥5× CpG/CpH coverage in ONT and ≥1× CpG/CpH coverage in both EM-seq and ACE-seq were used for mPFC regional comparisons. For single-site methylation level comparisons, only CpG/CpH sites with ≥10× coverage shared among ONT, EM-seq, and ACE-seq were included. To mitigate potential biases introduced by short-read mapping in repetitive regions, repeat elements across the genome were masked based on RepeatMasker annotations (https://www.repeatmasker.org) prior to global pairwise analyses. (CA)n repeats were specifically identified and profiled from RepeatMasker annotations for (CA)n repeat pairwise comparisons.

### False positive evaluation of ONT single-site modification calls

CpG sites with no detectable 5mCG signal, defined as EM-seq minus ACE-seq ≤ 0, were used to assess false positive patterns in ONT 5mCG calling, while sites with no detectable 5hmCG signal (ACE-seq = 0) were used to assess false positive patterns in ONT 5hmCG calling. Read strand information and the local 6-mer sequence context for each CpG site in the ONT data were profiled to evaluate strand-biased false positive 5hmCG calls. Representative false positive cases of ONT modification calls were visualized using Integrative Genomics Viewer (IGV, v2.17.4)^76^.

### 5hmC analysis across bacteria 5mC motifs

Modified base calling for bacteria samples was performed with DORADO with the genome reference for, *N. gonorrhoeae* FA1090 (RefSeq accession NC_002946.2), *H. pylori* strain JP26 (RefSeq accession NZ_CP023448.1) and *E. coli* K-12 MG1655 (RefSeq accession NC_000913.3). Cytosine sites within 5mC motif on bacteria genomes were retrieved using modkit with *modkit motif-bed* function. The motifs included GGCC and TCACC in *N. gonorrhoeae*, GCGC and GGCC in *H. pylori* and CCWGG in *E. coli*. For each motif, the counts of modified and unmodified bases from native and WGA samples were estimated using *modkit summary* with --no-sampling with adjusted P_mod_ (from 0.5 to 0.99) applied on *--filter-threshold*. The mean and standard deviation of 5hmC/C levels in native bacterial samples were then calculated across the five motifs.

### Ethics declarations

All animal procedures were conducted following protocols approved by the Rutgers University Institutional Animal Care and Use Committee (IACUC; protocol numbers 201900013 and 201900032). The kidney cancer samples used were under study IRB #19-00152 that was approved by the Institutional Review Board (IRB) of Mount Sinai. All tumor, blood and urine samples were collected with informed consent from participants in accordance with ethical guidelines. The participants were fully informed about the study’s aims and the voluntary nature of their participation, including the use of their samples for research purposes. Patient confidentiality was maintained throughout the study, and all procedures were conducted in compliance with relevant ethical standards. The postmortem brain samples from Alzheimer’s disease and healthy donors were exempt from IRB requirements, as determined by the Institutional Review Board (IRB) of Mount Sinai.

