## Supplementary Figures for "Incorporating FDR into the assessment of nanopore sequencing for the reliable detection of DNA modifications in real applications"

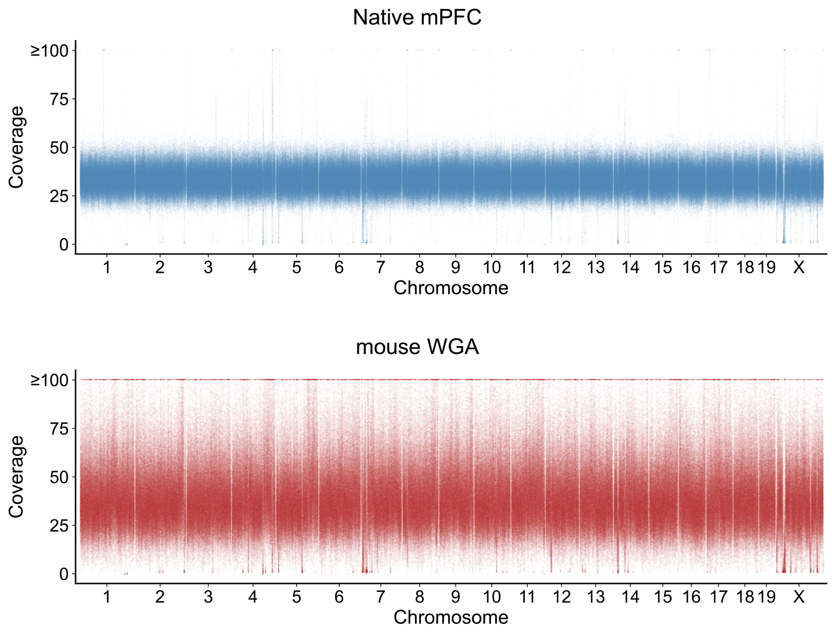


**Supplementary Figure. 1. Read coverage across all chromosomes for mPFC native and mouse WGA amplified samples.**

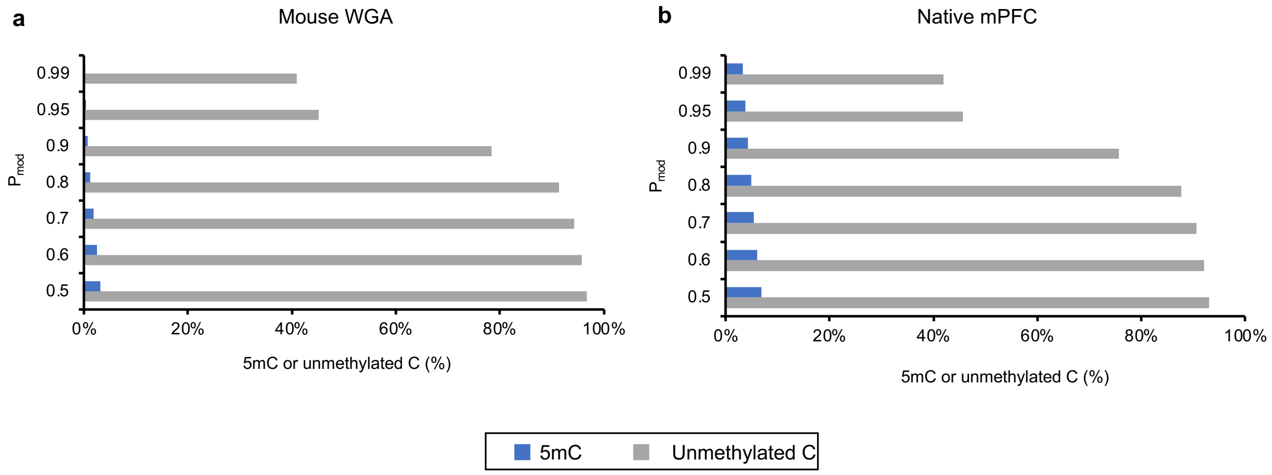


**Supplementary Figure 2. 5mC and unmethylated C levels in mouse WGA and native mPFC samples by DORADO 5mC model (v4.2.0) with different $P_{mod}$ threshold.**

(a) False positive 5mC or unmethylated C sites among the total C (%) in mPFC gDNA subjected to whole genome amplification (WGA). (b) 5mC or unmethylated C sites among the total C (%) in native mPFC gDNA. *x-axis*, 5mC or unmethylated C among the total C (%); *y-axis*, thresholds on modification probability ($P_{mod}$) with DORADO software. Only sites that successfully passed the $P_{mod}$ threshold (either confident unmodified or confident modified) were included for calculation.

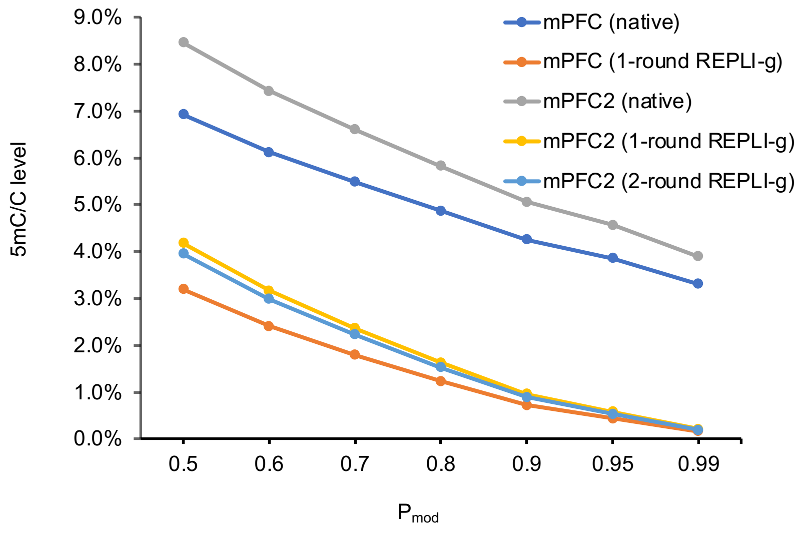


**Supplementary Figure 3. Levels of 5mC in native mPFC and mouse WGA samples prepared using different methods.**

mPFC (native) and mPFC2 (native from another female mouse, representing biological replicate) samples were each subjected to REPLI-g multiple displacement amplification to create mPFC (1-round REPLI-g) and mPFC2 (1-round REPLI-g) samples. mPFC2 (1-round REPLI-g) sample was subjected to a second round of REPLI-g amplification to create mPFC2 (2-round REPLI-g) sample. Results are based on DORADO 5mC model (v4.2.0). Refer to the Methods section for more details.

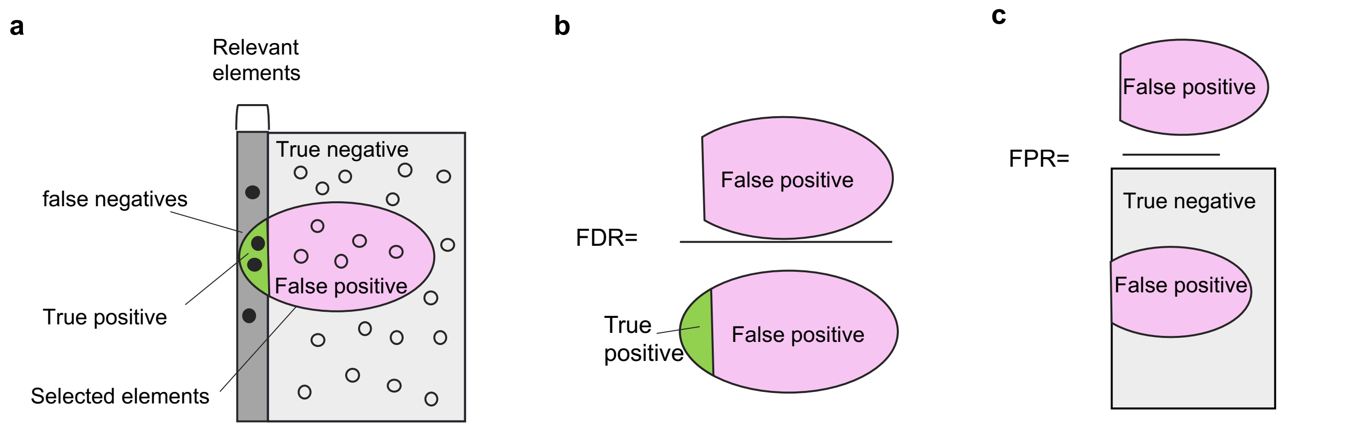


**Supplementary Figure 4. The use of FDR for modification evaluation when the modification level is low.**

(a) The left half of the image with the solid dots represents bases (e.g. A) that were modified with specific modification type (e.g. 6mA) across the genome, while the right half of the image with the hollow dots represents bases that were not modified (e.g. unmodified A) across the genome. The circle represents individual bases that were selected as modified.

(b) False discover rate (FDR) measures the proportion of false positives among all called positives.

(c) False positive rate (FPR) measures the proportion of false positives among all negative sites.

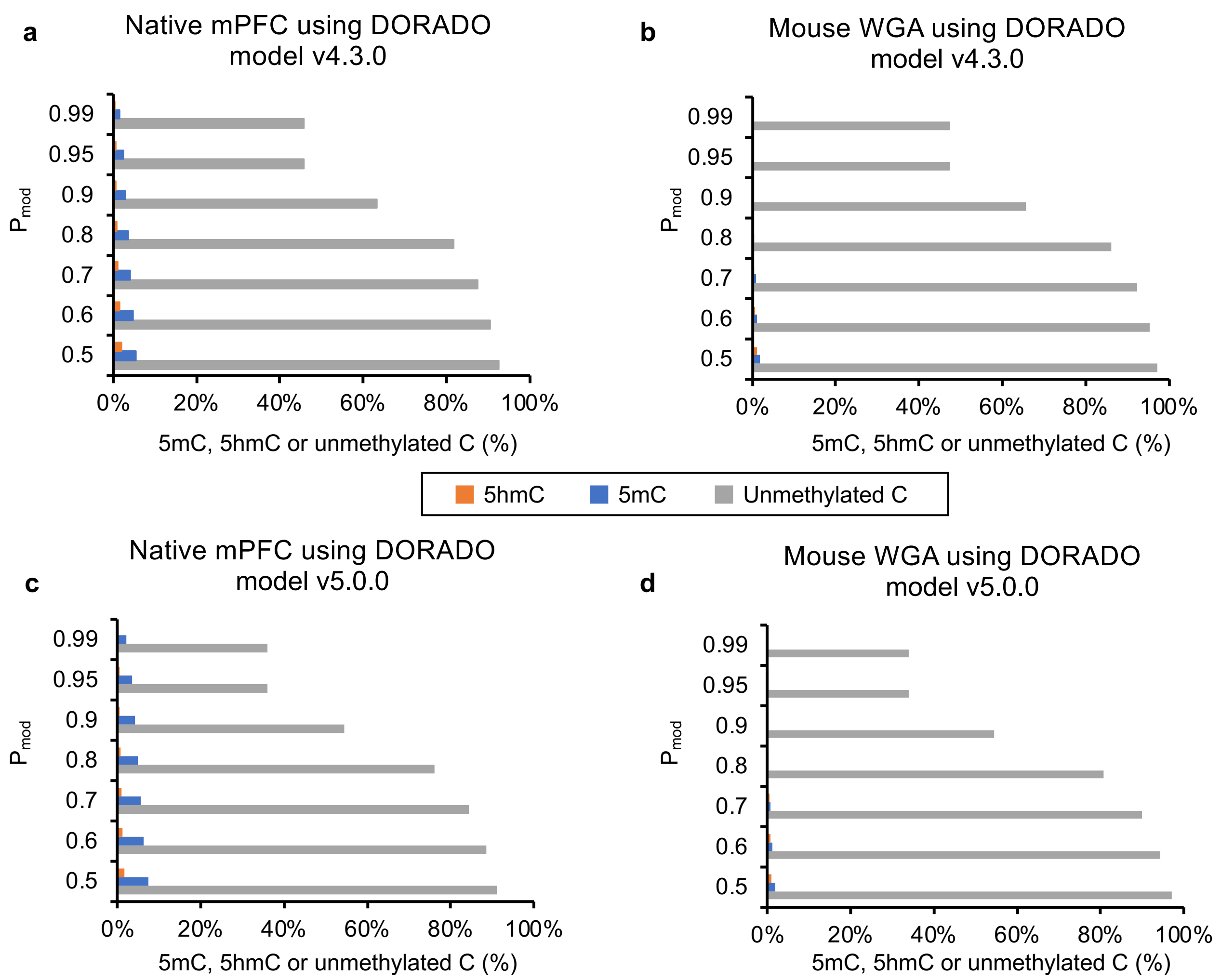


**Supplementary Figure 5. Levels of 5hmC, 5mC, and unmethylated C in native mPFC and mouse WGA samples using different versions of DORADO and base calling models.**

Proportions of 5hmC, 5mC, and unmethylated C are shown as a percentage of total cytosine sites. (**a**) and (**b**) display results for native mPFC gDNA and mouse WGA samples, respectively, using DORADO model v4.3.0. (**c**) and (**d**) show the corresponding results using DORADO model v5.0.0. Only sites that successfully passed the $P_{mod}$ threshold (either confident unmodified or confident modified) were included for calculation.

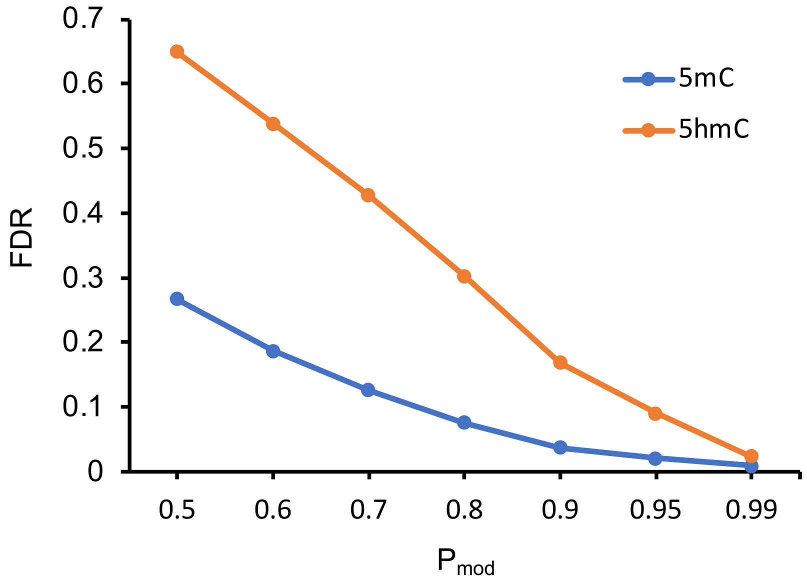


**Supplementary Figure 6. FDR evaluation of 5mC and 5hmC calls made from the native mPFC sample using DORADO 5mC_5hmC model v5.0.0.**

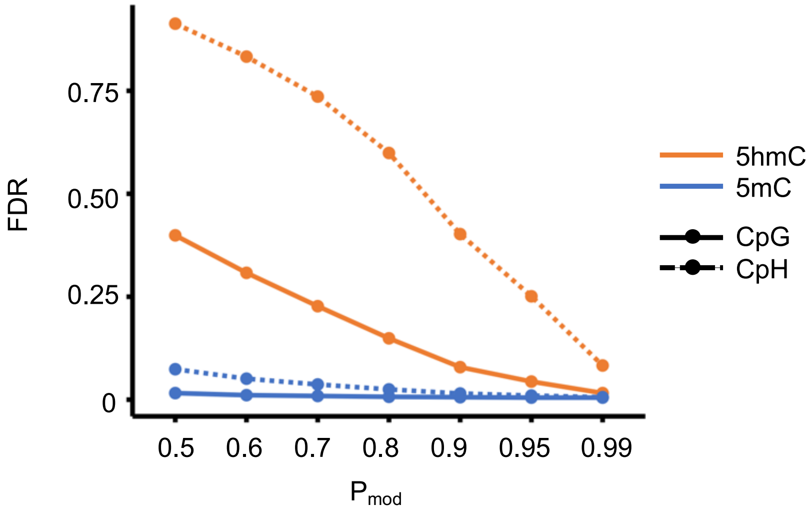


**Supplementary Figure 7. FDR evaluation of 5mC and 5hmC calls made at CpG and CpH sites in the native mPFC sample using DORADO 5mC_5hmC model v5.0.0.**

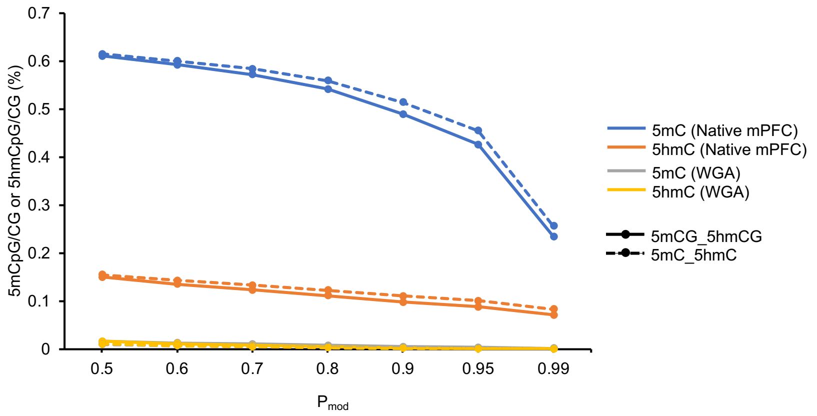


**Supplementary Figure 8. Read-level CpG modification analysis in the native mPFC and the mouse WGA detected with the 5mCG_5hmCG field vs the 5mC_5hmC field using DORADO model v4.3.0.**

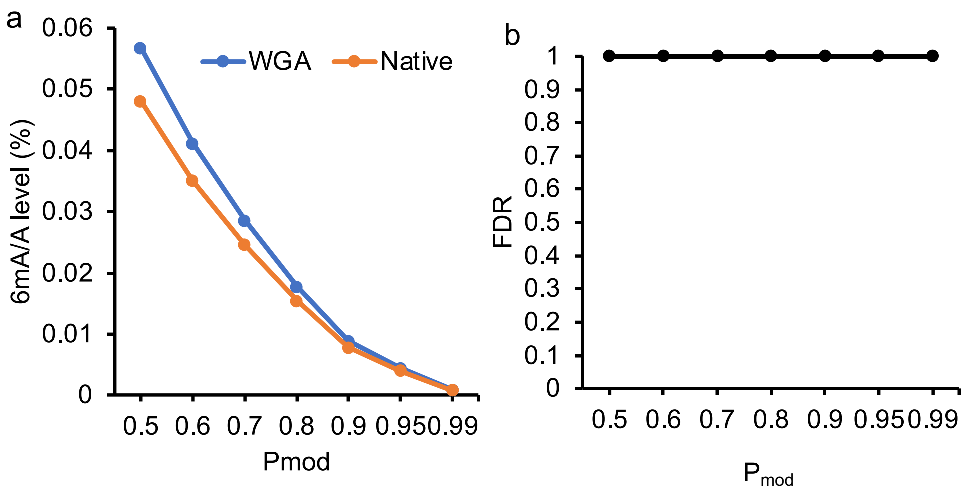


**Supplementary Figure 9. Read-level 6mA analysis in mPFC.**

(a) 6mA among the total A sites (%) in mouse WGA and the native mPFC gDNA by applying different thresholds on $P_{mod}$ using 6mA field in DORADO model v4.3.0.

(b) FDR evaluation for read-level modification analysis of 6mA in native mPFC.

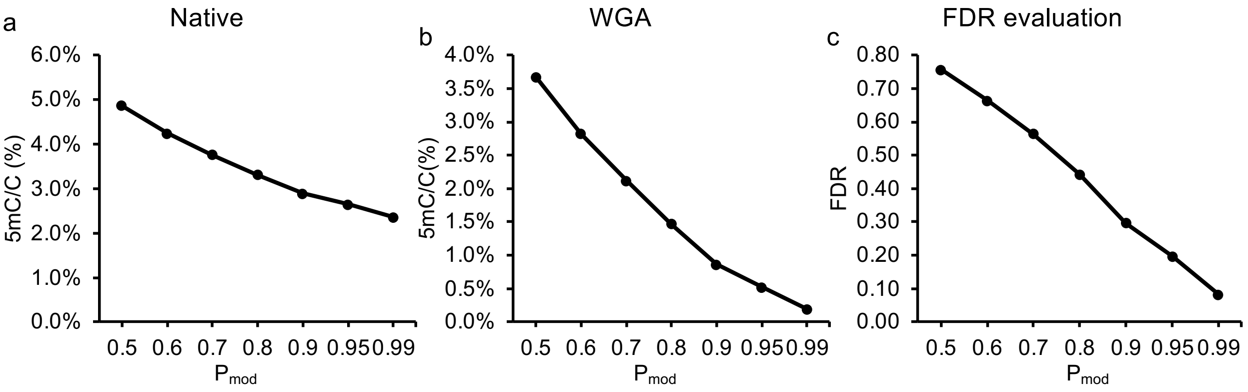


**Supplementary Figure 10. Read-level 5mC levels analysis in human Lymphoblastoid cell line (hLCL) samples by DORADO 5mC model v4.2.0 with different $P_{mod}$ threshold.**

(a) 5mC sites among the total C (%) in native hLCL gDNA. *x-axis*, thresholds on $P_{mod}$ by DORADO software.

(b) False positive 5mC sites among the total C (%) in hLCL gDNA subjected to WGA.

(c) FDR evaluation for read-level 5mC analysis in hLCL.

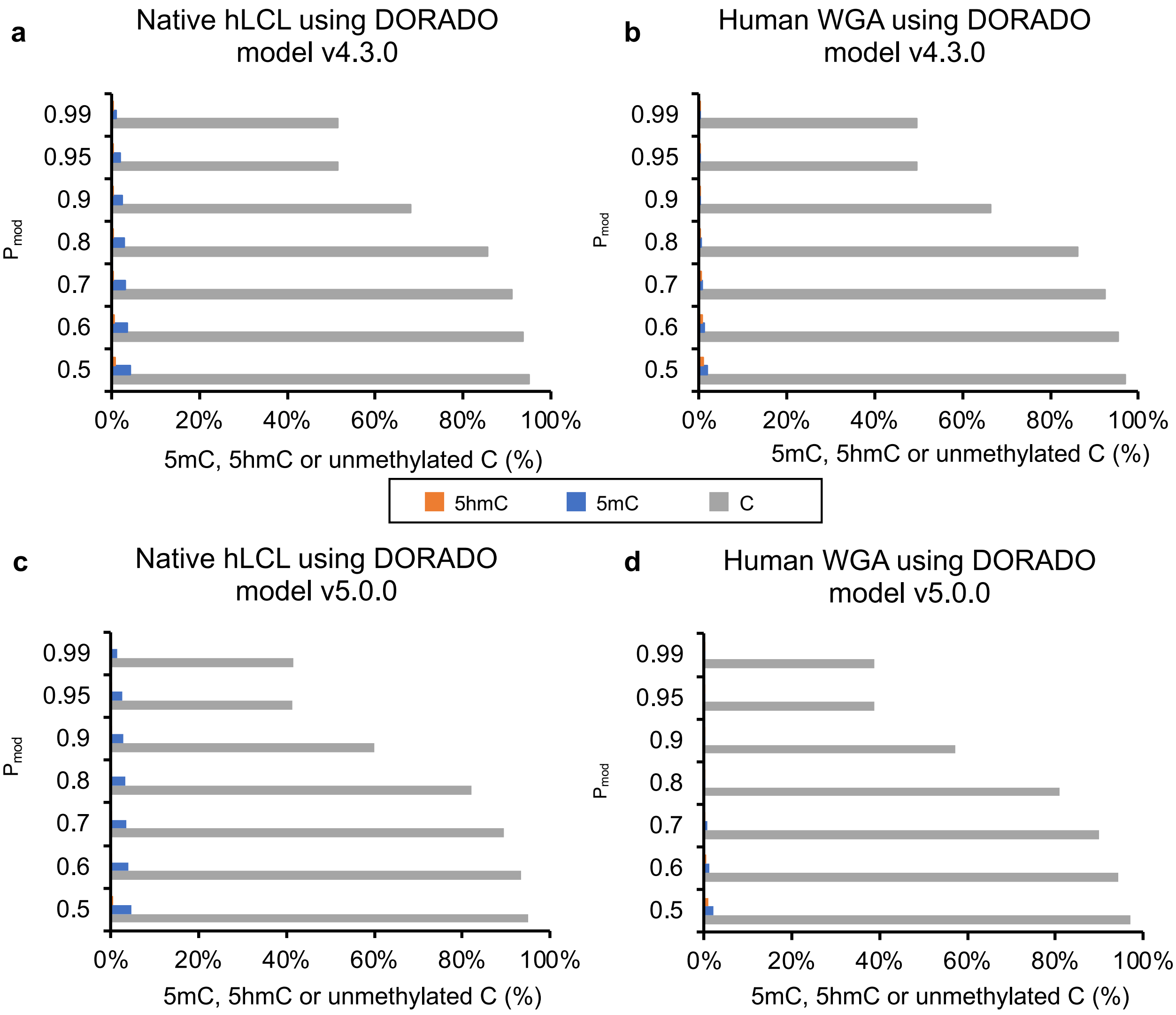


**Supplementary Figure 11. Levels of 5hmC, 5mC, and unmethylated C in native hLCL and human WGA samples using different DORADO base calling models.**

Proportions of 5hmC, 5mC, and unmethylated C are shown as a percentage of total cytosine sites. (**a**) and (**b**) display results for native hLCL gDNA and human WGA samples, respectively, using DORADO model v4.3.0. (**c**) and (**d**) show the corresponding results using DORADO model v5.0.0. Only sites that successfully passed the $P_{mod}$ threshold (either confident unmodified or confident modified) were included for calculation.

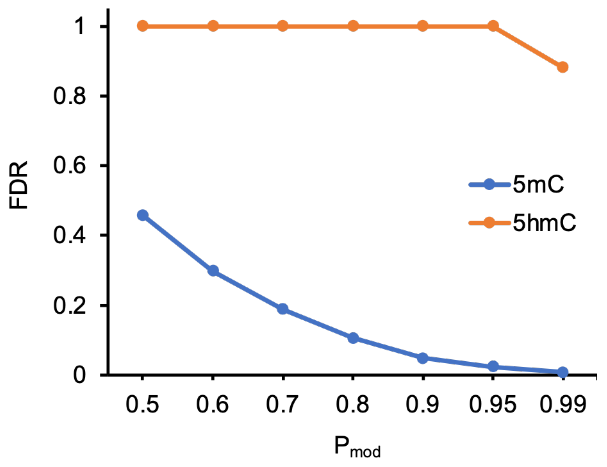


**Supplementary Figure 12. FDR evaluation of 5mC and 5hmC calls made from the native hLCL sample using DORADO 5mC_5hmC model v5.0.0.**

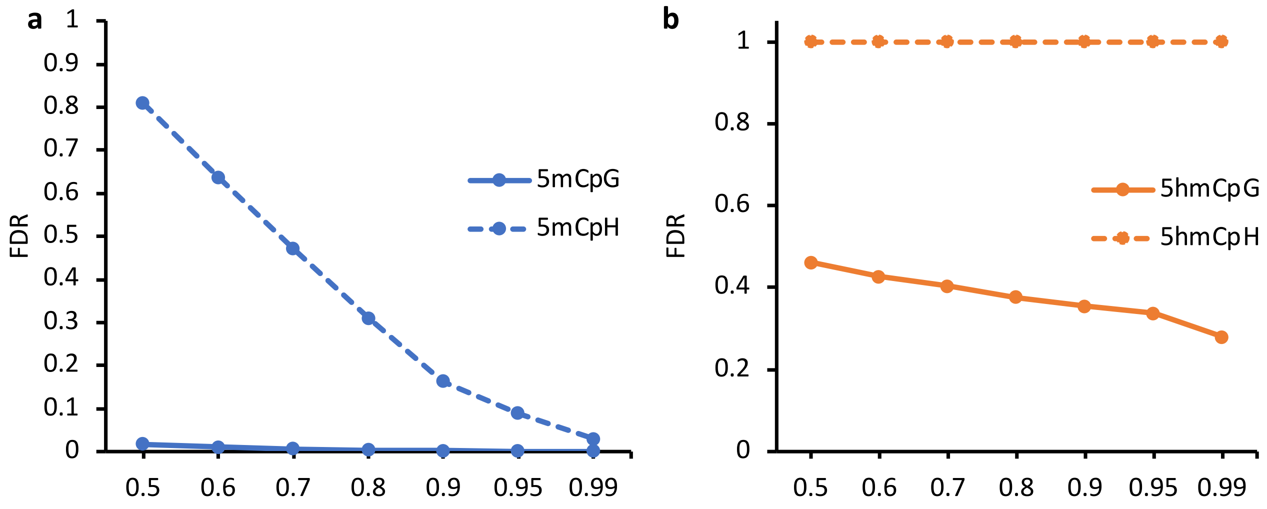


**Supplementary Figure 13. FDR evaluation of 5mC and 5hmC calls made at CpG and CpH sites in the native hLCL sample with DORADO 5mC_5hmC model v5.0.0.**

(**a**) FDR evaluation of 5mC calls made at CpG and CpH sites in the native hLCL sample. (**b**)FDR evaluation of 5hmC calls made at CpG and CpH sites in the native hLCL sample.

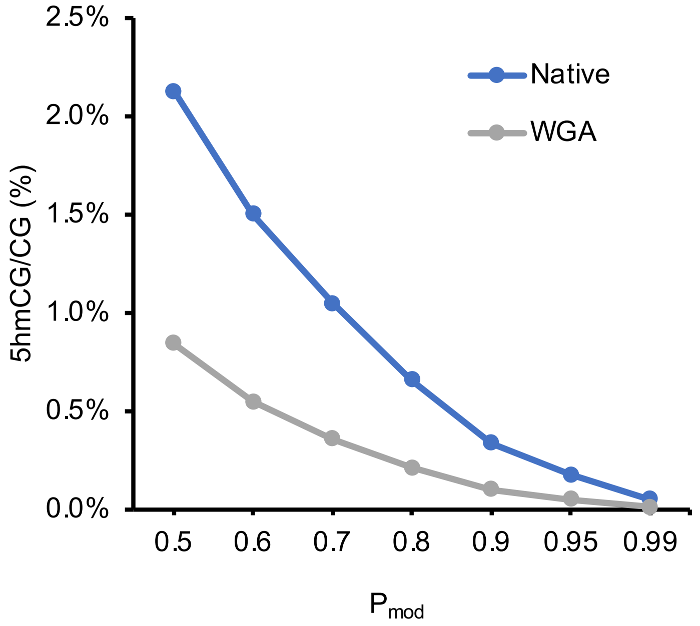


**Supplementary Figure 14. Read-level 5hmCG detected on hLCL Native and WGA gDNA.**

5hmCpG sites among the total CpG dinucleotide sites (%) in WGA and native hLCL gDNA by applying different thresholds on $P_{mod}$ using DORADO 5mC_5hmC model v4.3.0.

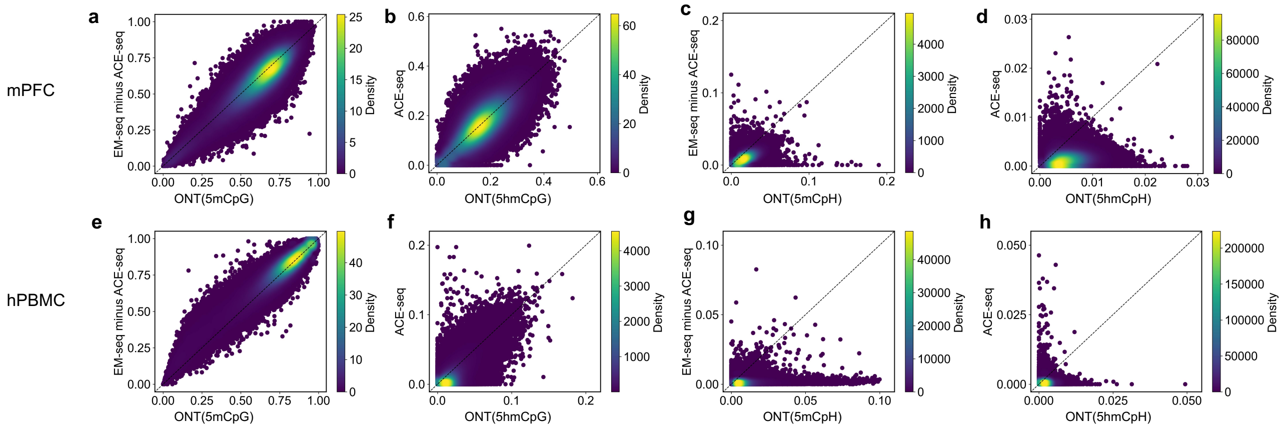


**Supplementary Figure 15. Comparison of overlapping 5mCpGs/5hmCpGs/5mCpHs/5hmCpHs between ONT and enzyme-based methods.**

Density plots of 5kb regions between ONT-based analysis (*x-axis*) and enzyme-based methods (*y-axis*) were shown for overlapped 5mCpGs, 5hmCpGs, 5mCpHs and 5hmCH in mPFC (a-d) and hPBMC (e-h). DORADO 5mC_5hmC model v4.3.0 and $P_{mod} \geq 0.75$ were used for ONT-based analyses.

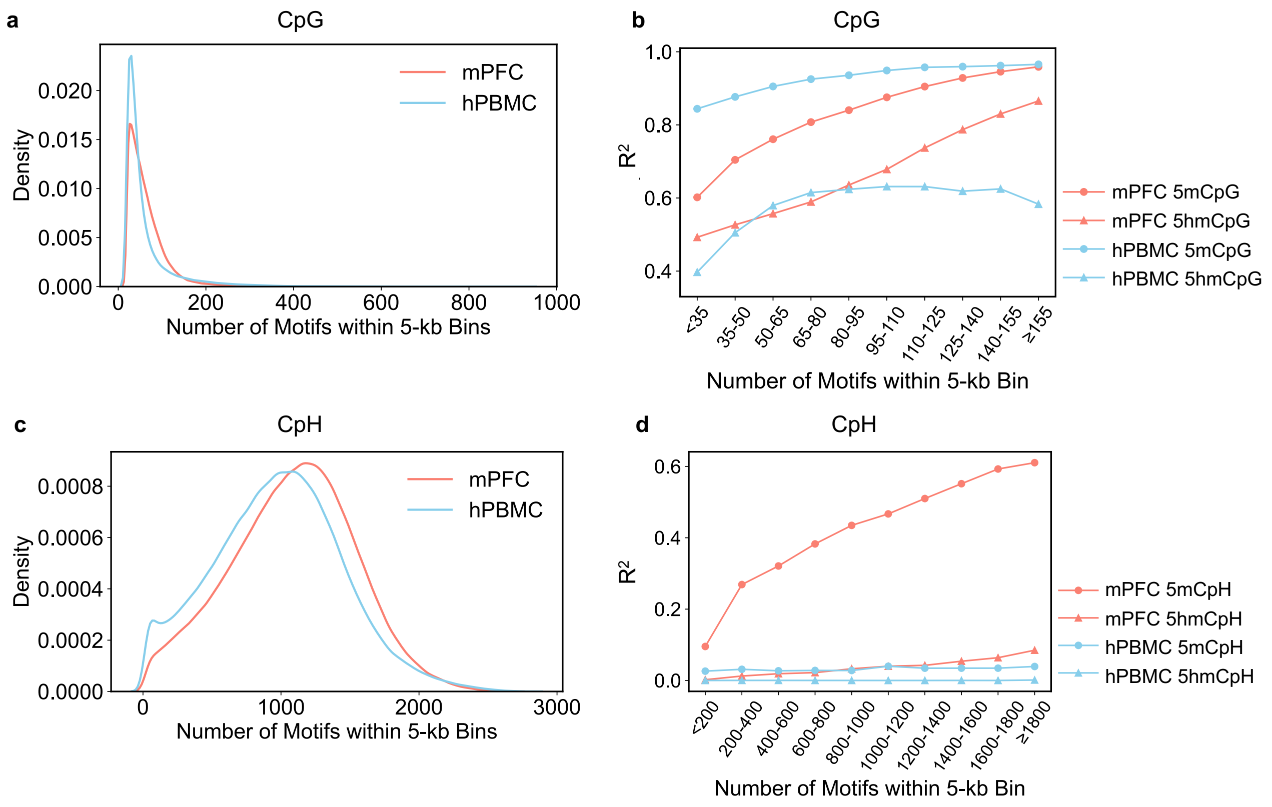


**Supplementary Figure 16. Comparison of ONT and EM-seq/ACE-seq for 5mC and 5hmC detection relative to motif density within 5kb regions.**

(a) Density plots of the distribution of CpG site counts within 5kb genomic windows for mPFC and hPBMC samples.

(b) Pearson correlation ($R^2$) between ONT and EM-seq/ACE-seq measurements for 5mC and 5hmC at CpG sites, stratified by CpG counts. DORADO 5mC_5hmC model v4.3.0 was used for ONT data analysis.

(c) Density plots of the distribution of CpH site counts within 5kb genomic windows for mPFC and hPBMC samples.

(d) Pearson correlation ($R^2$) between ONT and EM-seq/ACE-seq measurements for 5mC and 5hmC at CpH sites, stratified by CpG counts. DORADO 5mC_5hmC model v4.3.0 was used for ONT data analysis.

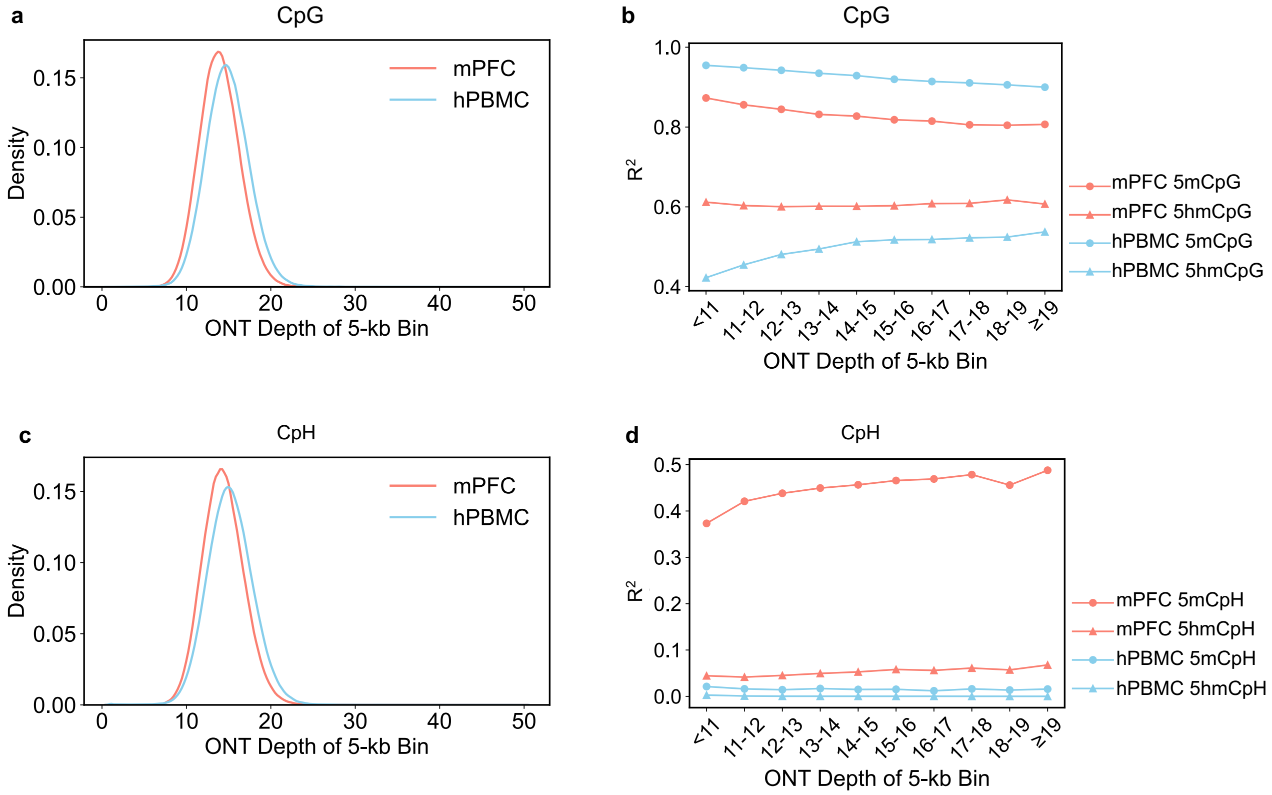


**Supplementary Figure 17. Comparison of ONT and EM-seq/ACE-seq for 5mC and 5hmC detection across different read coverage.**

(a) Density plots of the distribution of read depth for CpG sites within 5kb genomic windows for mPFC and hPBMC samples.

(b) Pearson correlation ($R^2$) between ONT and EM-seq/ACE-seq measurements for 5mC and 5hmC at CpG sites, stratified by read coverage. DORADO 5mC_5hmC model v4.3.0 was used for ONT data analysis.

(c) Density plots of the distribution of read depth for CpH sites within 5kb genomic windows for mPFC and hPBMC samples.

(d) Pearson correlation ($R^2$) between ONT and EM-seq/ACE-seq measurements for 5mC and 5hmC at CpG sites, stratified by read coverage. DORADO 5mC_5hmC model v4.3.0 was used for ONT data analysis.

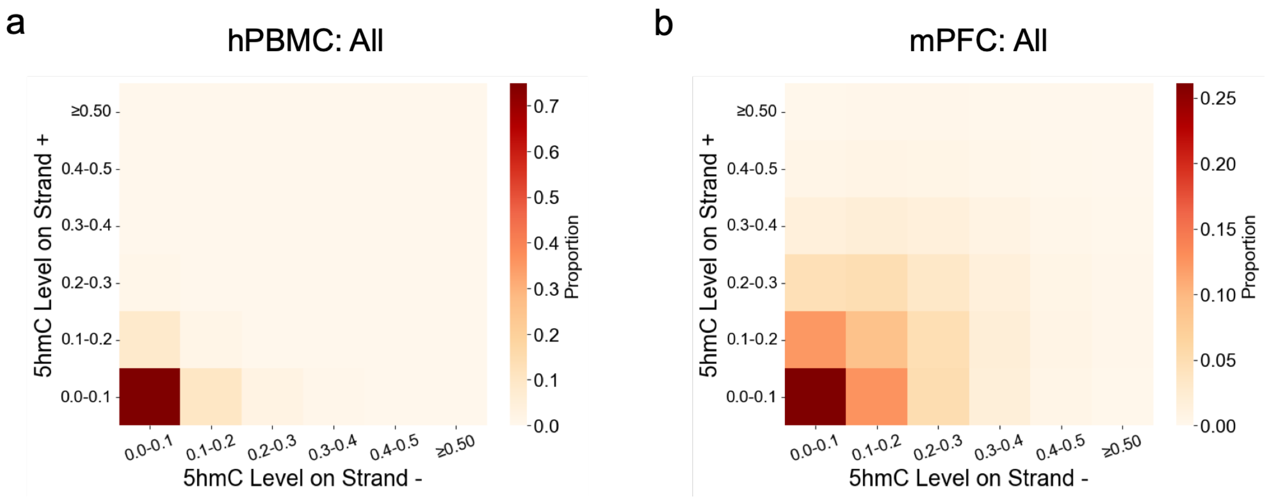


**Supplementary Figure 18. 5hmC levels across all genomic context in hPBMC and mPFC from ONT data.**

(a) Proportion of 5hmCG levels on both strands across hPBMC genome. *x-axis*, 5hmC level on strand -. *y-axis*, 5hmC level on strand +. False positive 5hmC sites were identified as CpG sites with 0% 5hmC levels in ACE-seq but ≥10% in ONT data. DORADO 5mC_5hmC model v4.3.0 was used for ONT data analysis.

(b) Equivalent analysis for 5hmCG on both strands at all CG loci across mPFC genome.

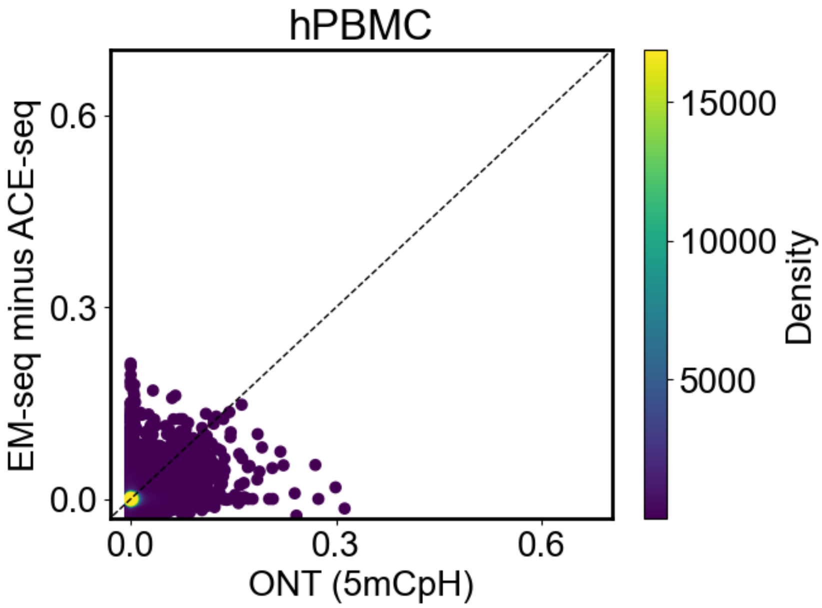


**Supplementary Figure 19. 5mCpH methylations on (CA)n repeats are not observed in hPBMC sample.**

Comparison of methylation level on (CA)n elements between EM-seq/ACE-seq and ONT in hPBMC sample. DORADO 5mC_5hmC model v4.3.0 was used for ONT data analysis.

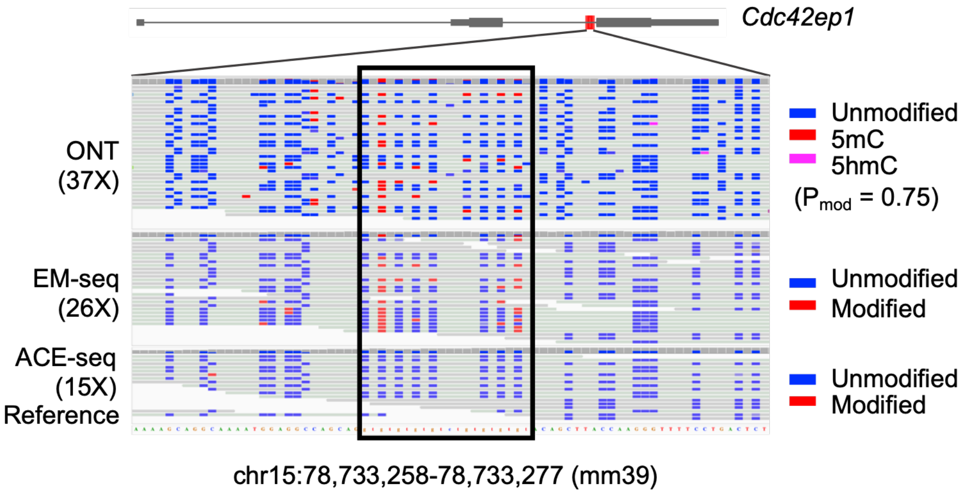


**Supplementary Figure 20. Example of 5mCpH methylation on a (CA)n repeat element located on the chr15 positive strand of mm39, consistently identified by EM-seq/ACE-seq and ONT methods.** Highlighted box indicates the region of (CA)n element. DORADO 5mC_5hmC model v4.3.0 was used for ONT data analysis. Regional sequencing coverage for ONT, EM-seq, and ACE-seq across the displayed IGV region is indicated in brackets.

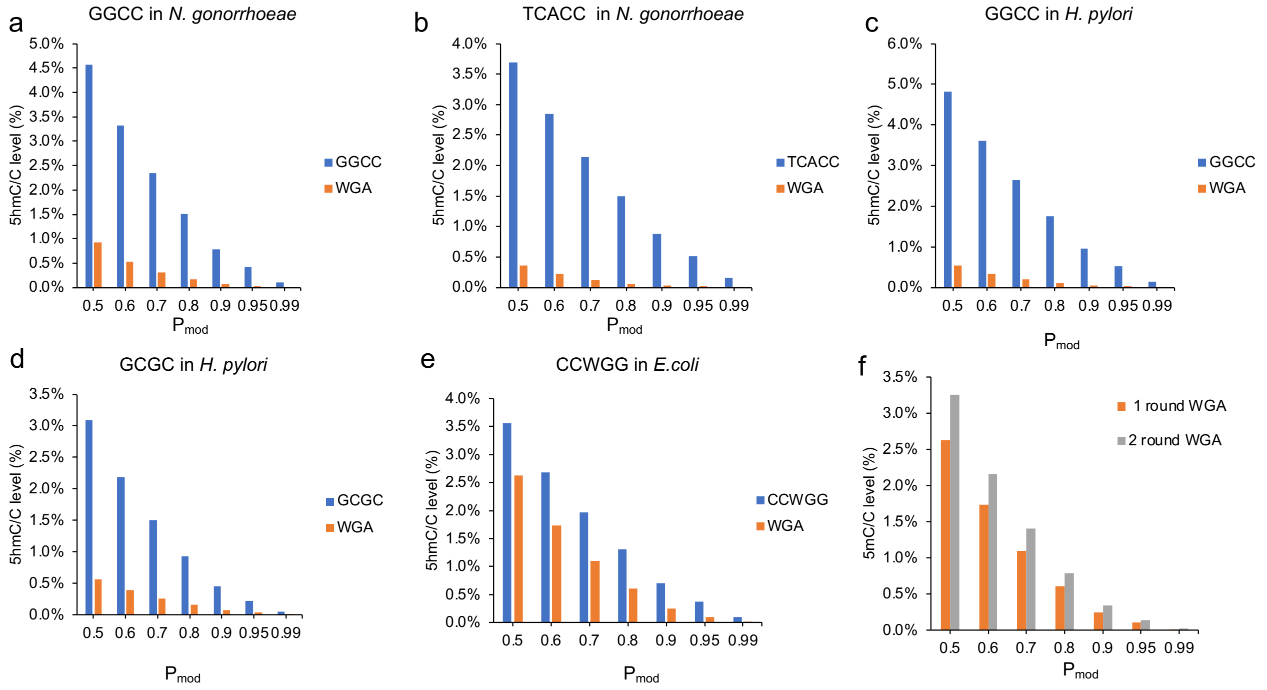


**Supplementary Figure 21. False positive 5hmC levels among 5mC-methylated motifs on nanopore reads from native bacterial genomes and matched WGA Samples.**

(a-e) 5hmC levels among 5mC sites (%) on reads from native bacterial gDNA samples were compared with those from the matched WGA samples for sites in motifs including: GGCC in *N. gonorrhoeae* (a), TCACC in *N. gonorrhoeae* (b), GGCC in *H. pylori* (c), GCGC in *H. pylori* (d), and CCWGG in *E. coli* (e). (f) An additional round of WGA using REPLI-g amplification on *E. coli* WGA samples confirmed reliable WGA amplification for *E. coli*. DORADO 5mC_5hmC model v4.3.0 was used for ONT data analyses.

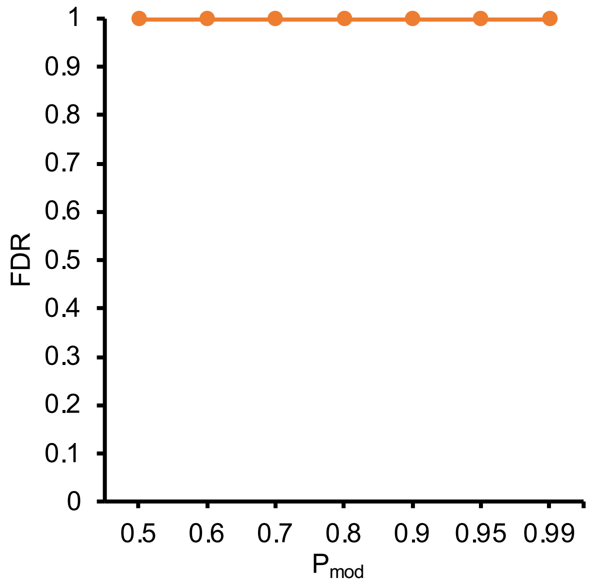


**Supplementary Figure 22. FDR evaluation for read-level 5hmC analysis in hLCL with additional bacteria 5mC motifs as the negative control. DORADO 5mC_5hmC model v4.3.0 was used for ONT data analysis.**

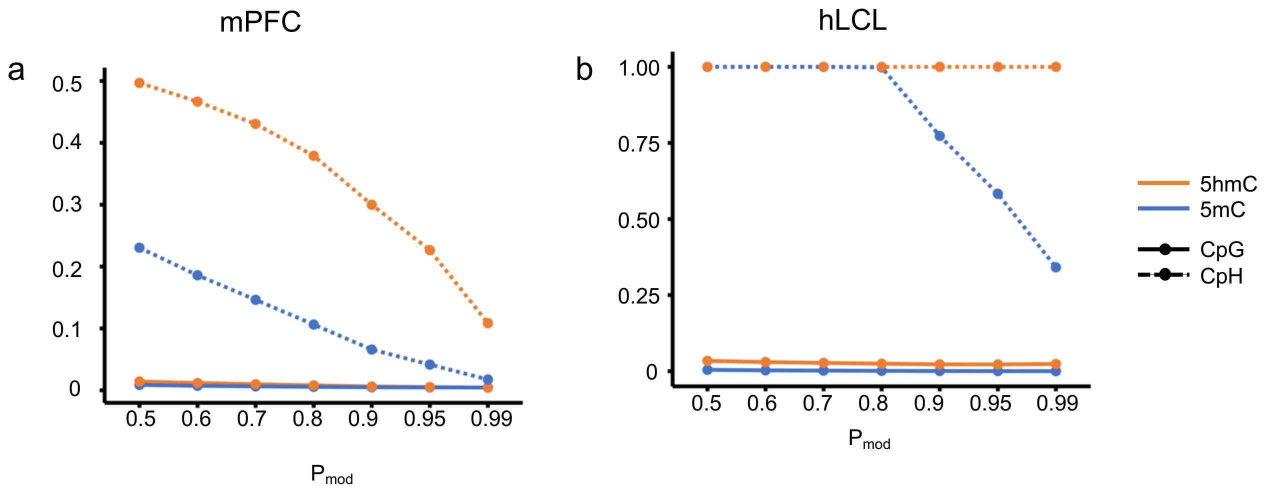


**Supplementary Figure 23. FDR evaluation of 5mC and 5hmC detection in mPFC and hLCL with dna_r10.4.1_e8.2_400bps_sup@v5.2.0@v1 model.**

FDR evaluation of 5mC and 5hmC calls made at CpG and CpH sites in the mPFC (a) and hLCL (b) samples.

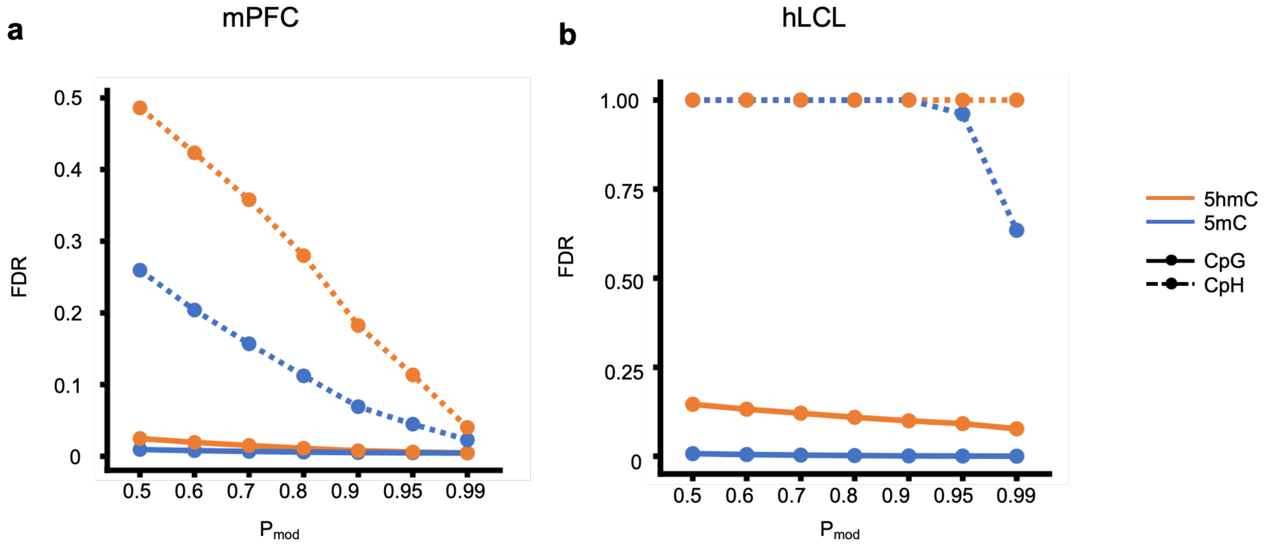


**Supplementary Figure 24. FDR evaluation of 5mC and 5hmC detection in mPFC and hLCL with dna_r10.4.1_e8.2_400bps_sup@v5.2.0@v2 model.**

FDR evaluation of 5mC and 5hmC calls made at CpG and CpH sites in the mPFC (a) and hLCL (b) samples.

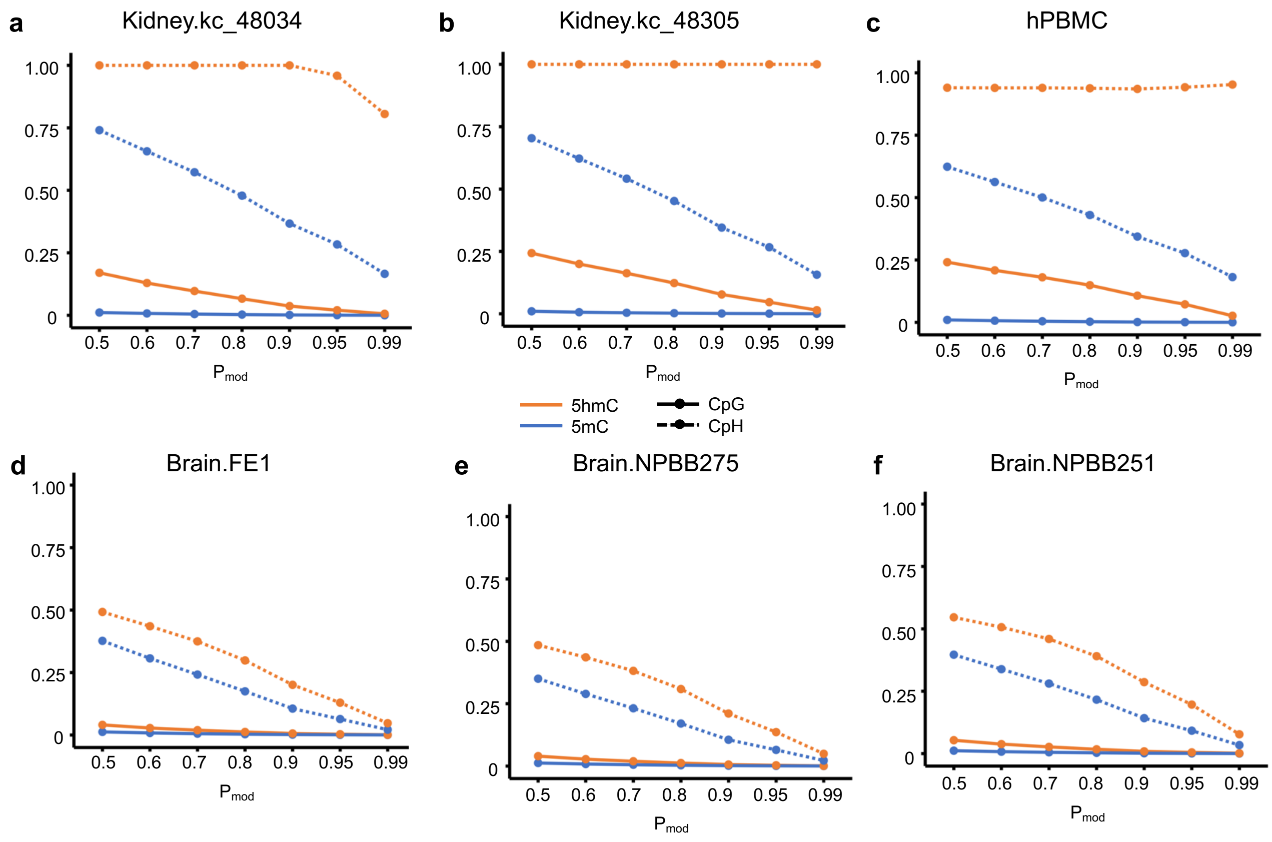


**Supplementary Figure 25. 5mC and 5hmC detection in multiple human tissues calls analyzed with DORADO model v4.3.0.**

FDR evaluation of 5mC and 5hmC calls made at CpG and CpH sites in the two human kidney tissues (a-b), hPBMC (c) and three human brain tissues (c-e).

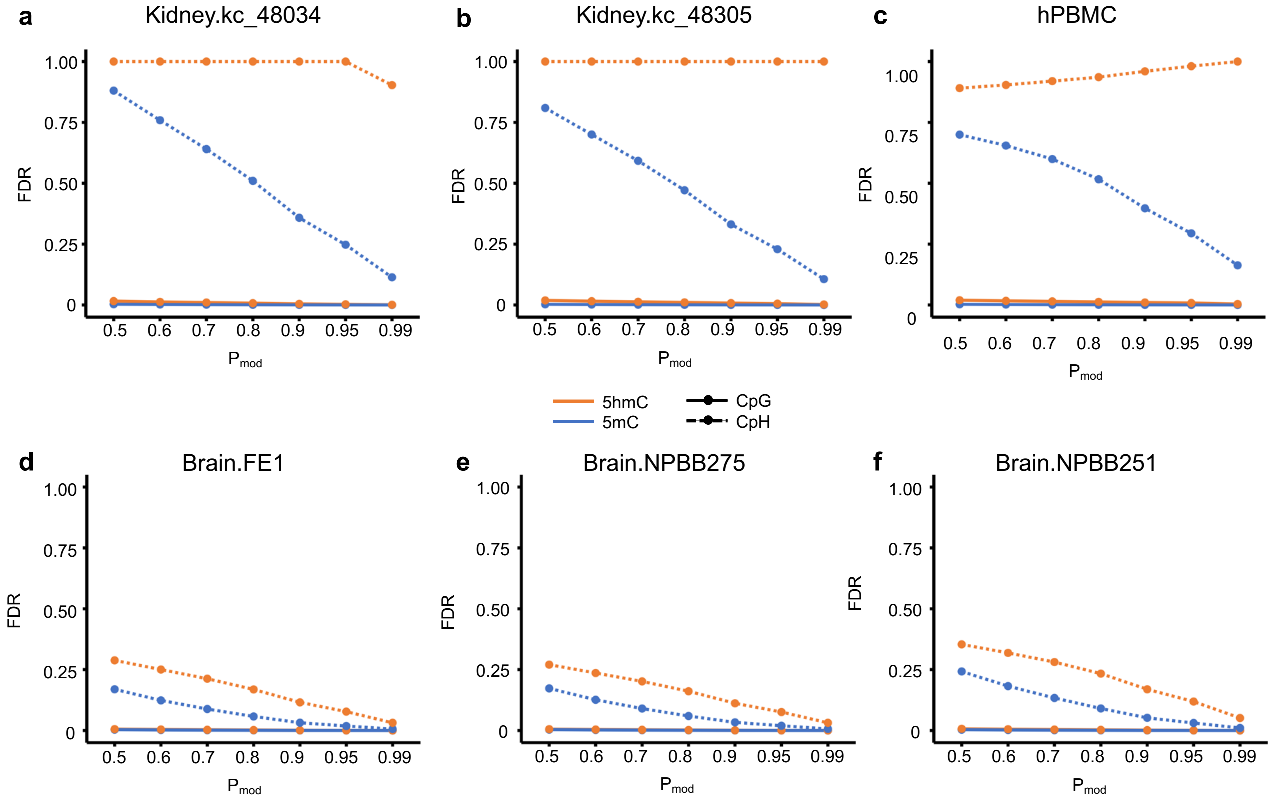


**Supplementary Figure 26. 5mC and 5hmC detection in multiple human tissues calls analyzed with DORADO model v5.2.0@v1.**

FDR evaluation of 5mC and 5hmC calls made at CpG and CpH sites in the two human kidney tissues (a-b), hPBMC (c) and three human brain tissues (c-e).

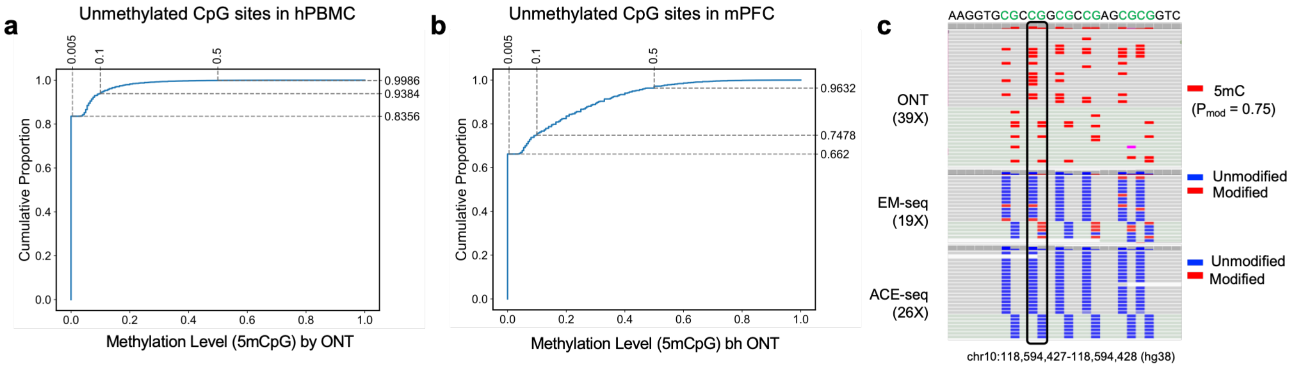


**Supplementary Figure 27. False-positive 5mCpG calls analyzed with DORADO base calling model v5.2.0@v1.**

(a) False-positive 5mCpG calls based on gold standard independent methods (EM-seq/ACE-seq) in hPBMC. Cumulative proportion of CpG sites called by DORADO at loci with 0% methylation (unmethylated CpG sites) based on EM-seq/ACE-seq. Bins (0.005) show cumulative fractions; x-axis, ONT methylation levels; y-axis: cumulative proportion of loci below each threshold.

(b) Equivalent analysis for mPFC.

(c) Example of recurrent false-positive 5mCpG calls by ONT (hPBMC), not supported by EM-seq/ACE-seq, visualized in IGV. Regional sequencing coverage for ONT, EM-seq, and ACE-seq across the displayed IGV region is indicated in brackets.

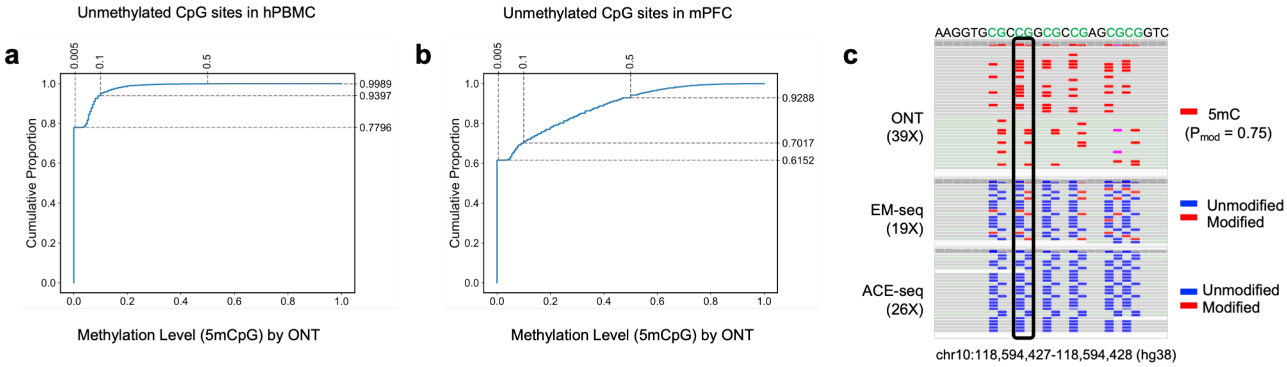


**Supplementary Figure 28. False-positive 5mCpG calls analyzed with DORADO base calling model v5.2.0@v2.**

(a) False-positive 5mCpG calls based on gold standard independent methods (EM-seq/ACE-seq) in hPBMC. Cumulative proportion of CpG sites called by DORADO at loci with 0% methylation (unmethylated CpG sites) based on EM-seq/ACE-seq. Bins (0.005) show cumulative fractions; x-axis, ONT methylation levels; y-axis: cumulative proportion of loci below each threshold.

(b) Equivalent analysis for mPFC.

(c) Example of recurrent false-positive 5mCpG calls by ONT (hPBMC), not supported by EM-seq/ACE-seq, visualized in IGV. Regional sequencing coverage for ONT, EM-seq, and ACE-seq across the displayed IGV region is indicated in brackets.

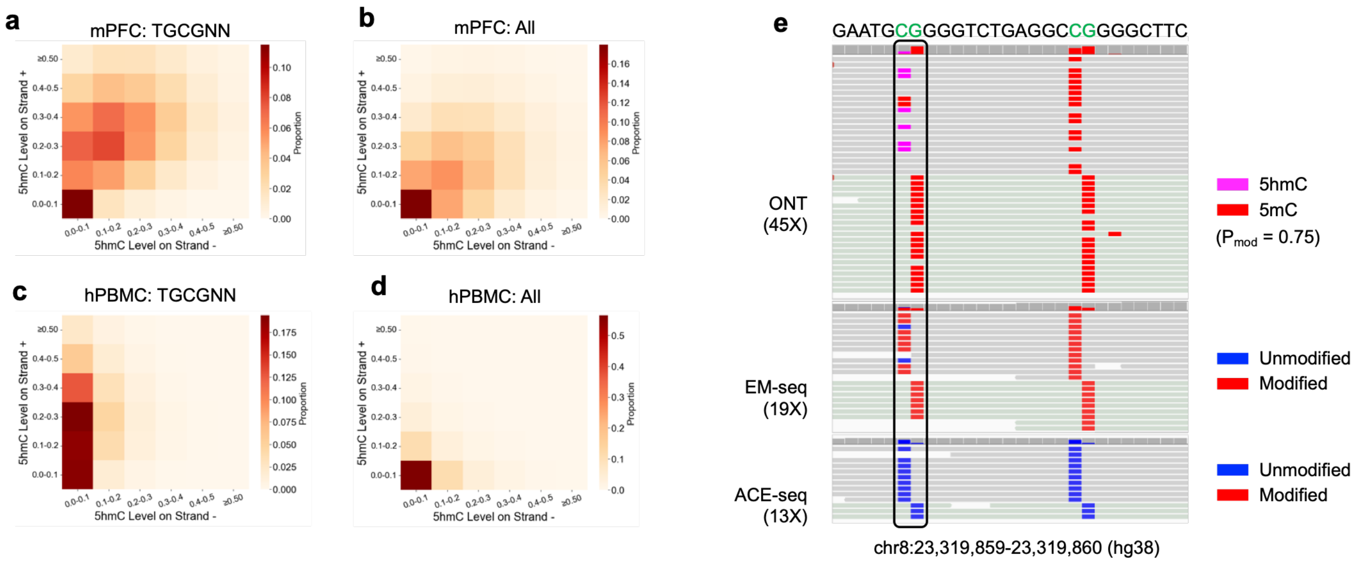


**Supplementary Figure 29. Strand-biased false positive 5hmC calls under the TGCGNN context from ONT data analyzed with DORADO base calling model v5.2.0@v1.**

(a) Proportion of 5hmCG on both strands at TGCGNN loci across mPFC genome. *x-axis*, 5hmC level on strand -. *y-axis*, 5hmC level on strand +. False positive 5hmC sites were identified as CpG sites with 0% 5hmC levels in ACE-seq but ≥10% in ONT data.

(b) Equivalent analysis for 5hmCG on both strands at all CG loci across mPFC genome.

(c) Equivalent analysis for 5hmCG on both strands at TGCGNN loci across hPBMC genome.

(d) Equivalent analysis for 5hmCG on both strands at CG loci across hPBMC genome.

(e) Example of repeatedly false positive ONT 5hmCG calls with a hemi-methylation pattern in the hPBMC sample. Regional sequencing coverage for ONT, EM-seq, and ACE-seq across the displayed IGV region is indicated in brackets.

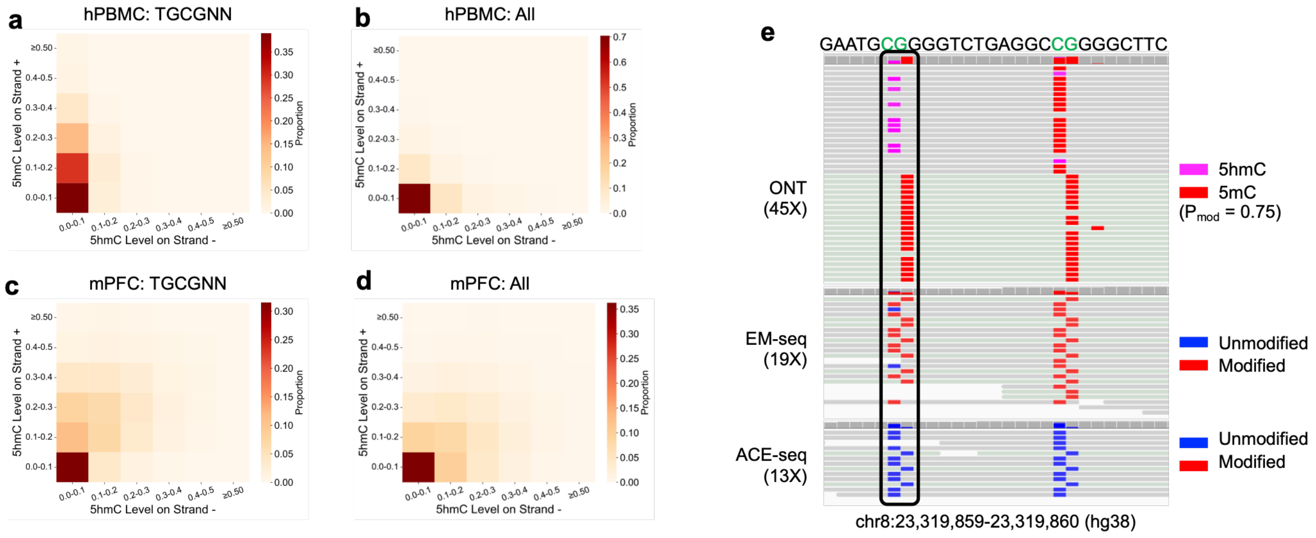


**Supplementary Figure 30. Strand-biased false positive 5hmC calls under the TGCGNN context from ONT data analyzed with DORADO base calling model v5.2.0@v2.**

(a) Proportion of 5hmCG on both strands at TGCGNN loci across mPFC genome. x-axis, 5hmC level on strand -. y-axis, 5hmC level on strand +. False positive 5hmC sites were identified as CpG sites with 0% 5hmC levels in ACE-seq but ≥10% in ONT data.

(b) Equivalent analysis for 5hmCG on both strands at all CG loci across mPFC genome.

(c) Equivalent analysis for 5hmCG on both strands at TGCGNN loci across hPBMC genome.

(d) Equivalent analysis for 5hmCG on both strands at all CG loci across hPBMC genome.

(e) Example of repeatedly false positive ONT 5hmCG calls with a hemi-methylation pattern in the hPBMC sample. Regional sequencing coverage for ONT, EM-seq, and ACE-seq across the displayed IGV region is indicated in brackets.

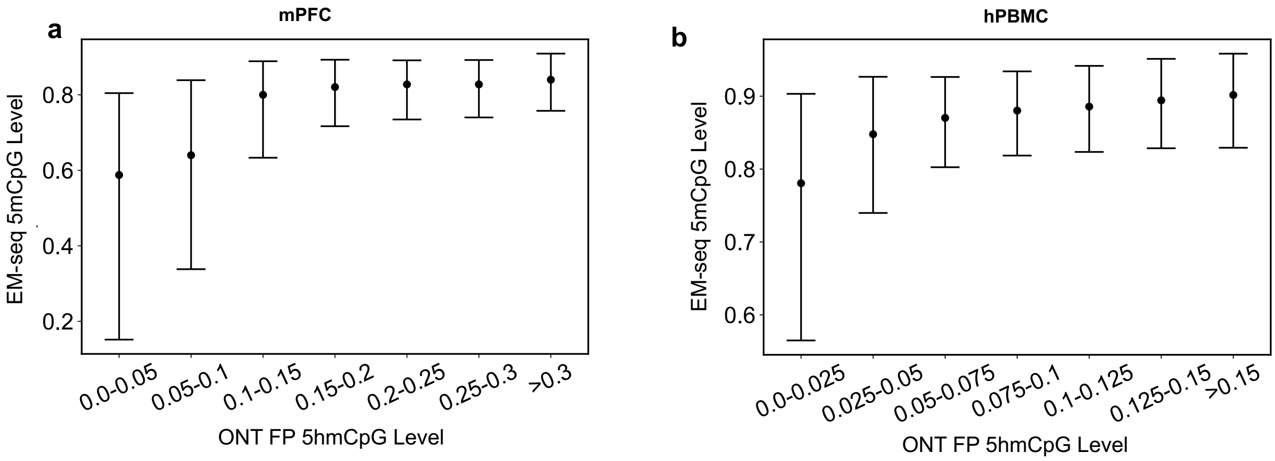


**Supplementary Figure 31. False positive 5hmCpG by confounding 5mCpG analyzed with DORADO model v5.2.0@v1.**

(a) 5mCpG levels by EM-seq across different false positive (FP) 5hmCpG levels in ONT data across mPFC genome.

(b) Equivalent analysis for hPBMC genome.

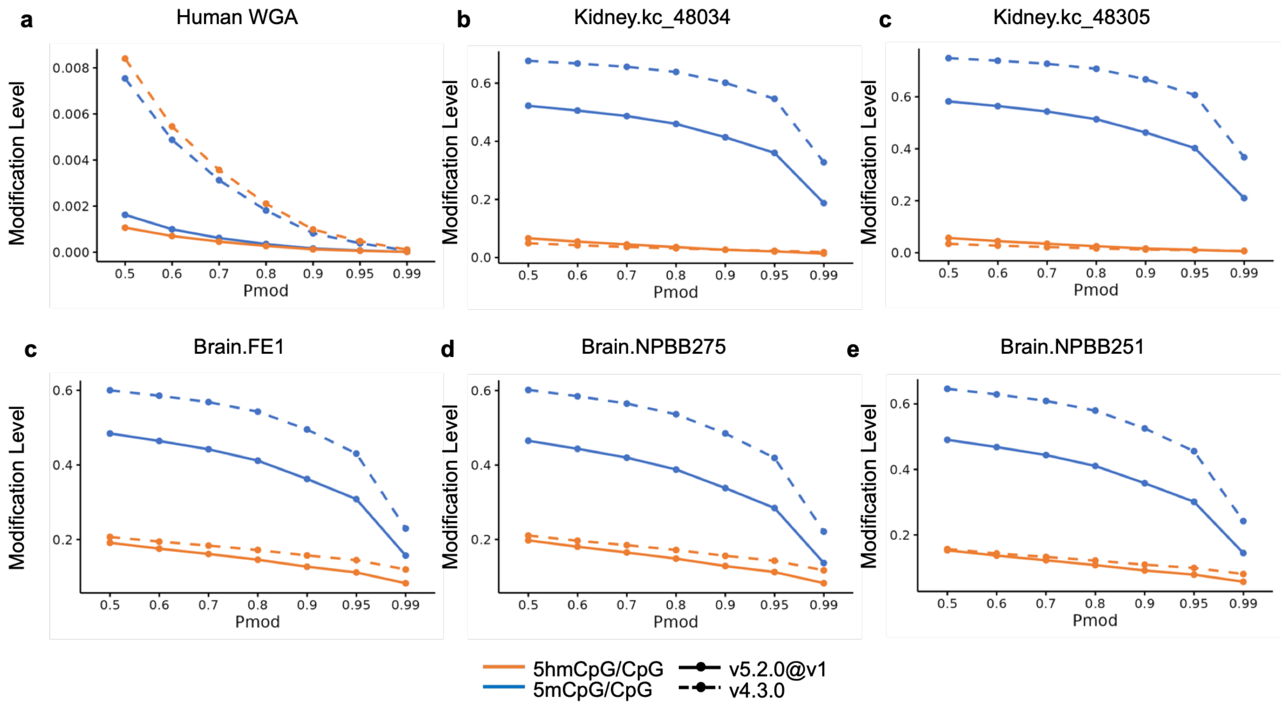


**Supplementary Figure 32. Modification levels of 5mCpG and 5hmCpG among all CpG sites in multiple human tissues with different DORADO models.**

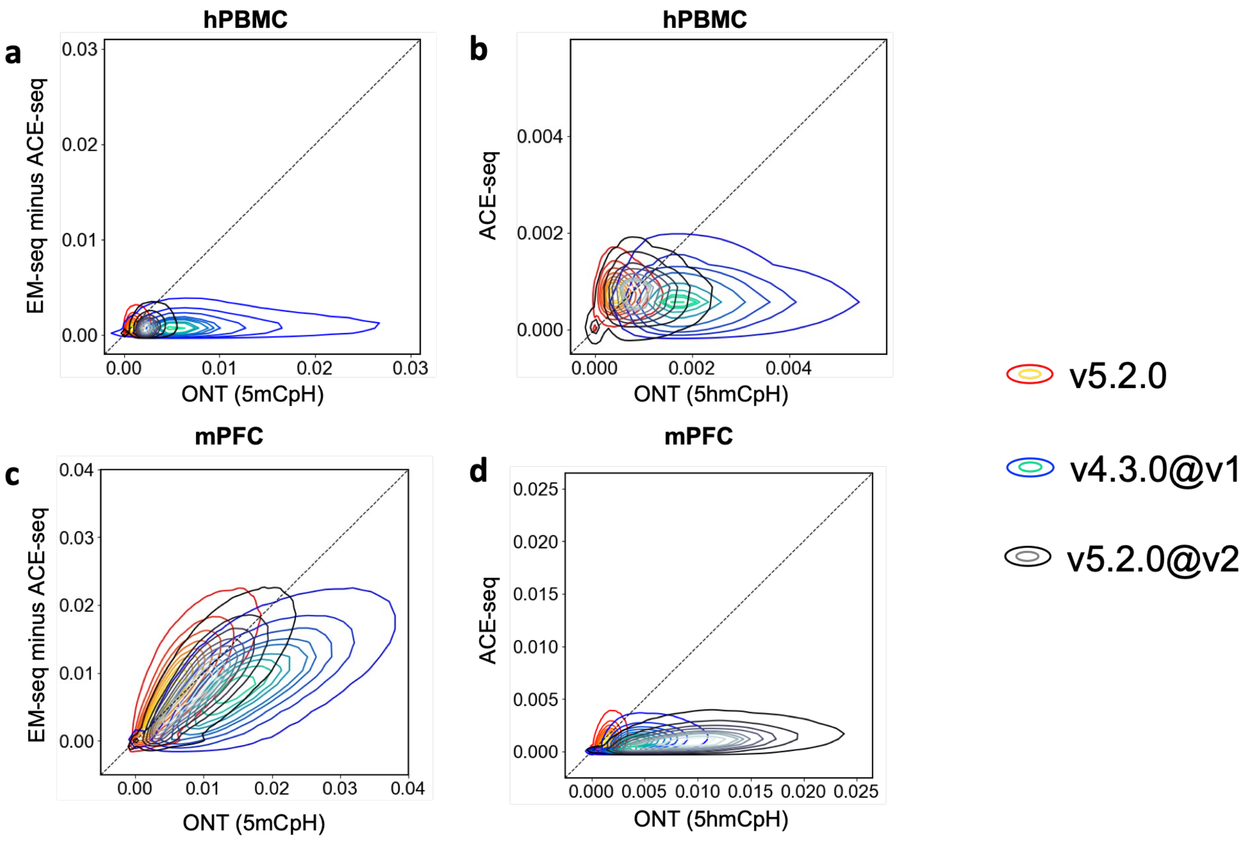


**Supplementary Figure 33. Comparison of overlapping CpHs between ONT analyzed with different DORADO base calling models and enzyme-based methods.**

Contour plots of CpH levels within overlapped 5kb regions between ONT-based analysis (*x-axis*), analyzed by DORADO models v4.3.0, v5.2.0@v1 and v5.2.0@v2, and enzyme-based methods (*y-axis*) for 5mCpHs mPFC (a), 5hmCpH in mPFC (b), 5mCpHs in hPBMC (c) and 5hmCpH in hPBMC (d). $P_{mod} \geq 0.75$ was used for ONT-based analyses.

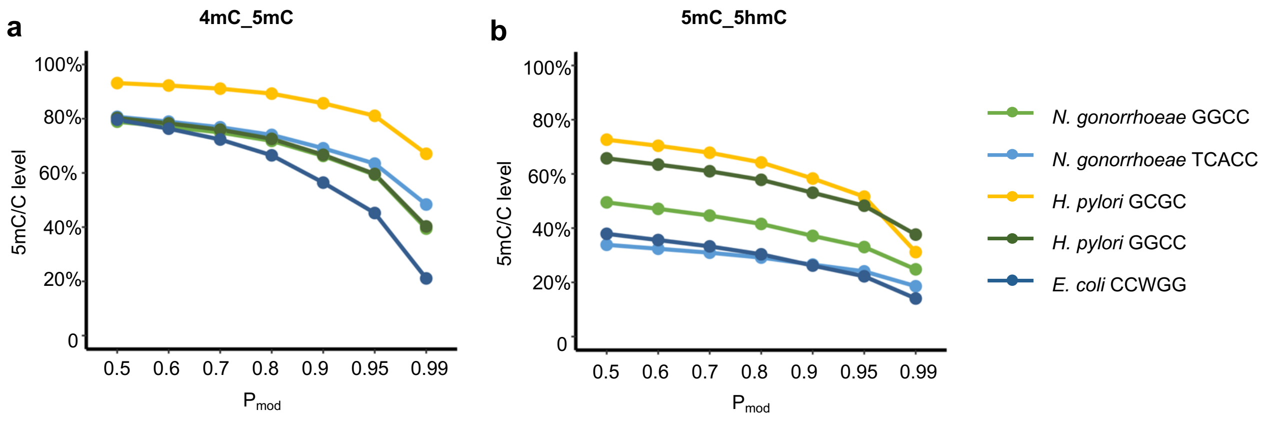


**Supplementary Figure 34. 5mC/C level within 5mC motifs in bacteria using DORADO v5.2.0@v1 model with 5mC_5hmC and 4mC_5mC fields.**

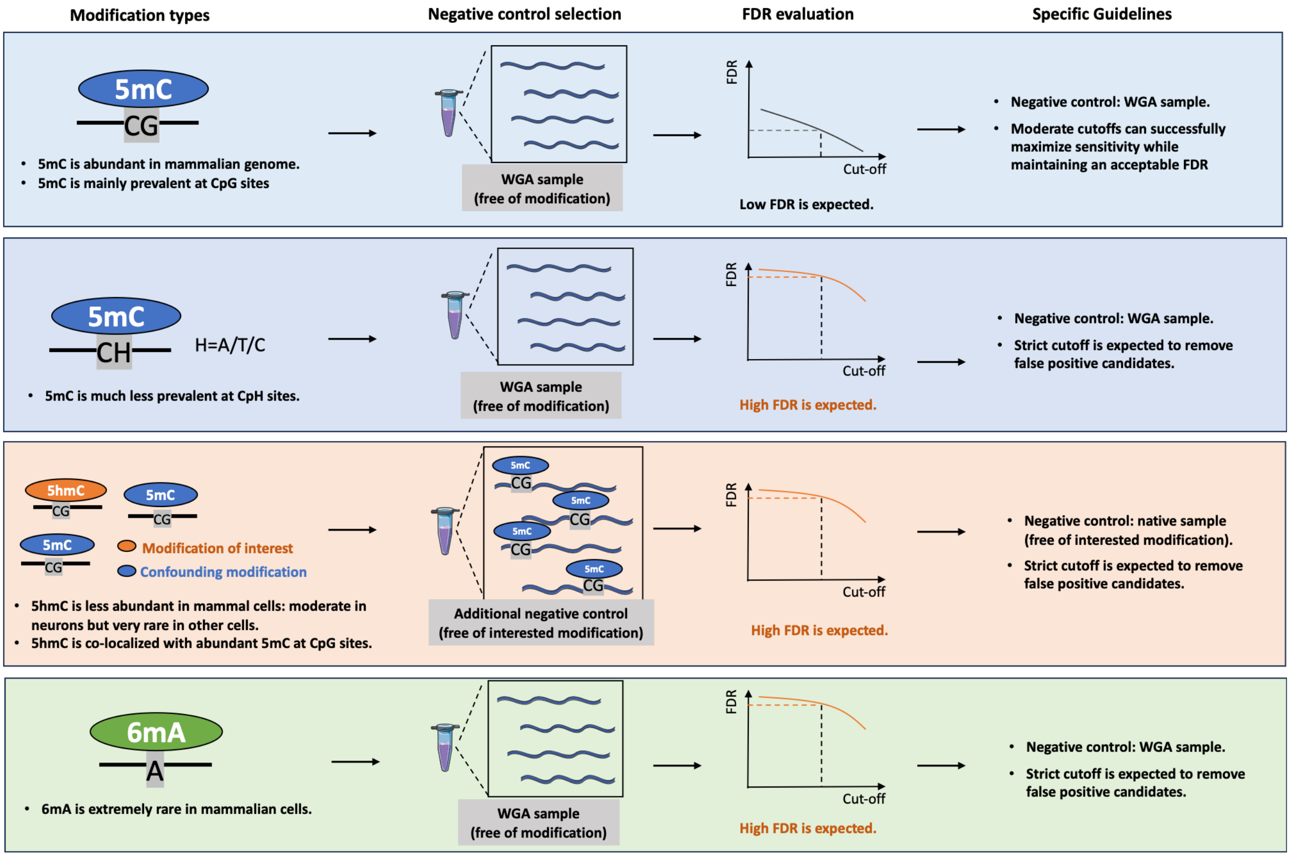


**Supplementary Figure 35. Specific guideline for modification detections in mammalian cells using modFDR.**

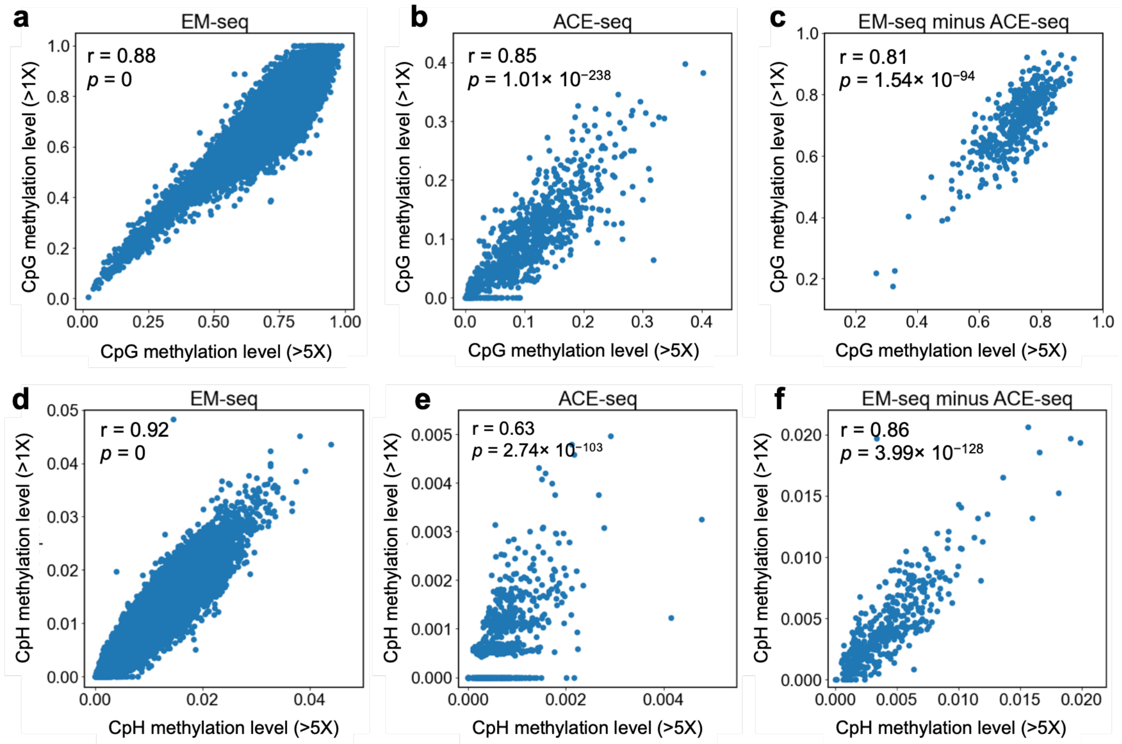


**Supplementary Figure 36. High correlated regional methylation levels between high and low coverage for CpG and CpH sites in mPFC.**

Regions from mPFC EM-seq and ACE-seq with high coverage (>5×) were subsampled to simulate low-coverage (>1×) conditions. Pearson correlation coefficients were calculated to compare CpG and CpH methylation levels between the original high-coverage and subsampled low-coverage data for EM-seq, ACE-seq, and the difference between them (EM-seq minus ACE-seq).
